# Extraordinary peptide-binding mode of a songbird MHC class-I molecule suggests mechanism to counter pathogen immune evasion

**DOI:** 10.1101/2023.03.13.532050

**Authors:** Sandra Eltschkner, Samantha Mellinger, Sören Buus, Morten Nielsen, Kajsa M Paulsson, Karin Lindkvist-Petersson, Helena Westerdahl

## Abstract

Long-distance migratory animals such as birds and bats have evolved to withstand selection imposed by pathogens across the globe, and pathogen richness is known to be particularly high in tropical regions. Immune genes, so-called Major Histocompatibility Complex (MHC) genes, are highly duplicated in songbirds compared to other vertebrates, and this high MHC diversity has been hypothesised to result in a unique adaptive immunity. To understand the rationale behind the evolution of the high MHC genetic diversity in songbirds, we determined the structural properties of an MHC class I protein, Acar3, from a long-distance migratory songbird, the great reed warbler *Acrocephalus arundinaceus* (in short: *Acar*). The structure of Acar3 was studied in complex with pathogen-derived antigens and shows an overall antigen presentation similar to human MHC class I. However, the peptides bound to Acar3 display an unusual conformation: Whereas the N-terminal ends of the peptides display enhanced flexibility, the conformation of their C-terminal halves is rather static. This uncommon peptide-binding mode in Acar3 is facilitated by a central Arg residue within the peptide-binding groove that fixes the backbone of the peptide at its central position, and potentially permits successful interactions between MHC class I and innate immune receptors. Our study highlights the importance of investigating the immune system of wild animals, such as birds and bats, to uncover unique immune mechanisms which may neither exist in humans nor in model organisms.

## Introduction

Songbirds belong to the most species-rich bird order on earth, Passeriformes [1]. They cross the globe during their annual migratory journeys and manage to breed in a wide range of different habitats [2-4]. The successful radiation with regards to number of species suggests that songbirds have been very adaptable to different environments, and hence able to handle selection from a wide range of pathogens [5-7]. However, these adaptations are not unique to songbirds, as all vertebrates in the tropics, where pathogens are particularly numerous, have evolved immune systems capable of withstanding a diverse spectrum of pathogens [8-10]. In recent time, the immune system in bats, forming the second largest order among mammals, has been scrutinised in detail since many bats carry a multitude of viruses (Ebola, Nipah, severe acute respiratory syndrome (SARS) and Middle East respiratory syndrome (MERS)) seemingly without any symptoms, whereas humans become severely ill upon infection with these viruses [11-16]. It is less known to what extent songbirds carry pathogens without showing symptoms yet bearing the risk of becoming zoonotic.

Interestingly, bats and songbirds display a similar evolutionary adaptation in their adaptive immune system, since their Major Histocompatibility Complex class I (MHC-I) genes have expanded more than in other terrestrial vertebrates, with many songbirds harbouring vastly duplicated MHC-I genes (> 50 genes per individual) [17-20]. MHC-I molecules are expressed on all nucleated cells and are central in every adaptive immune response towards intracellular pathogens, such as viruses and intracellular bacteria [21]. MHC-I molecules present antigens to CD8^+^ T-cells, and if the antigen is pathogen-derived, the cell will be killed, whereas cells only displaying self-antigens are left untouched. Each MHC-I molecule can present a small fraction of antigens. The range of those antigens, which contain a preferred set of anchor residues, is determined by the properties of the peptide-binding groove (PBG).

Theoretically, a very high MHC diversity (number of different MHC alleles per individual) is unfavourable as MHC diversity higher than the optimal diversity will diminish the total T-cell repertoire [22]. However, the strength of the negative selection of T-cells, *i*.*e*. central tolerance, is correlated not only with the MHC diversity *per se*, but also with the average antigen-binding spectrum of individual MHC-I molecules: a narrow repertoire allows a higher MHC diversity than a broader repertoire per MHC-I molecule [23, 24]. Humans have six MHC-I genes, three classical and three non-classical, whereas in bats up to 12 MHC-I-like genes have been reported [17] which corresponds well with bats having a potentially narrower peptide-binding repertoire per individual MHC-I molecule than humans [25-27]. Songbirds have at least 20 expressed MHC-I alleles [28], *i*.*e*. between 10 and 20 MHC-I genes depending on degree of heterozygosity, suggesting that songbird MHC-I molecules could have an even more restricted repertoire than humans and bats. Although the average antigen-binding breadth is expected to differ among species, MHC-I molecules with narrow (fastidious) and broad (promiscuous) antigen binding repertoires probably exists in most species, as shown in recent comparative work from humans and the avian model species, chicken (*Gallus gallus*), the latter with only two classical MHC-I genes [29].

Here we set out to determine the structural properties of Acar3, the most highly expressed MHC-I molecule from a long-distant migratory songbird, the great reed warbler *Acrocephalus arundinaceus* (*Acar*). In this species, that breeds in the Western Palearctic and winters in Sub-Saharan Africa [30], the MHC-I diversity has been thoroughly characterised [31-33]. Our structural studies on the Acar3-heavy chain in complex with *Acar* beta-2-microglobulin (β_2_m) and two different peptides reveal a surprisingly high N-terminal malleability of those peptides, despite sharing a Met residue acting as N-terminal anchor. Intriguingly, an Arg residue at the centre of the PBG which binds to the backbone of the peptides leads to highly similar conformations within their C-terminal halves. This unique peptide conformation in Acar3 could have important implications for the immune competence of the great reed warbler, potentially enabling a more effective control of infection through counteracting pathogen escape mechanisms.

## Results

### The top-binding peptides of Acar3 show a preference for distinct anchor residues

MHC-I molecules bind short peptides of varying lengths, and after testing the length preferences of two different great reed warbler MHC-I molecules, (***Figure S1***), we applied a 9-mer positional scanning combinatorial peptide library (PSCPL) [34] to select peptides that can be presented by Acar3. The PSCPL analysis revealed two dominant anchors in Acar3, at peptide positions three and nine, with a clear preference for hydrophobic residues, in particular for Met (M) at position three, and Phe (F) at position nine ***(Figure 1, Figure S2)***. Moreover, there was a tendency towards hydrophobic residues in the middle of the peptides (positions five and six) such as Ala (A), and Pro (P). At position two, a preference for Met (M) could be observed, but also for other amino acids with long and flexible side chains, such as Lys (K) and Arg (R). The stability of Acar3 in complex with 94 different peptides indicated as suitable binders by the PSCPL analysis was measured using a Scintillation Proximity Assay (SPA). From this assay, the three top-ranked peptides with respect to half-lives of Acar3 complexes, *i*.*e*. peptide 1 (AMSAQAAAF, “P1”; T½ = 11.3 h), peptide 2 (YMTLQAVTF, “P2”; T½ = 11.2 h) and peptide 3 (MTMITPPTF; “P3”; T½ = 10.3 h) ***(Table S1)***, were selected for further studies.

**Figure 1:**
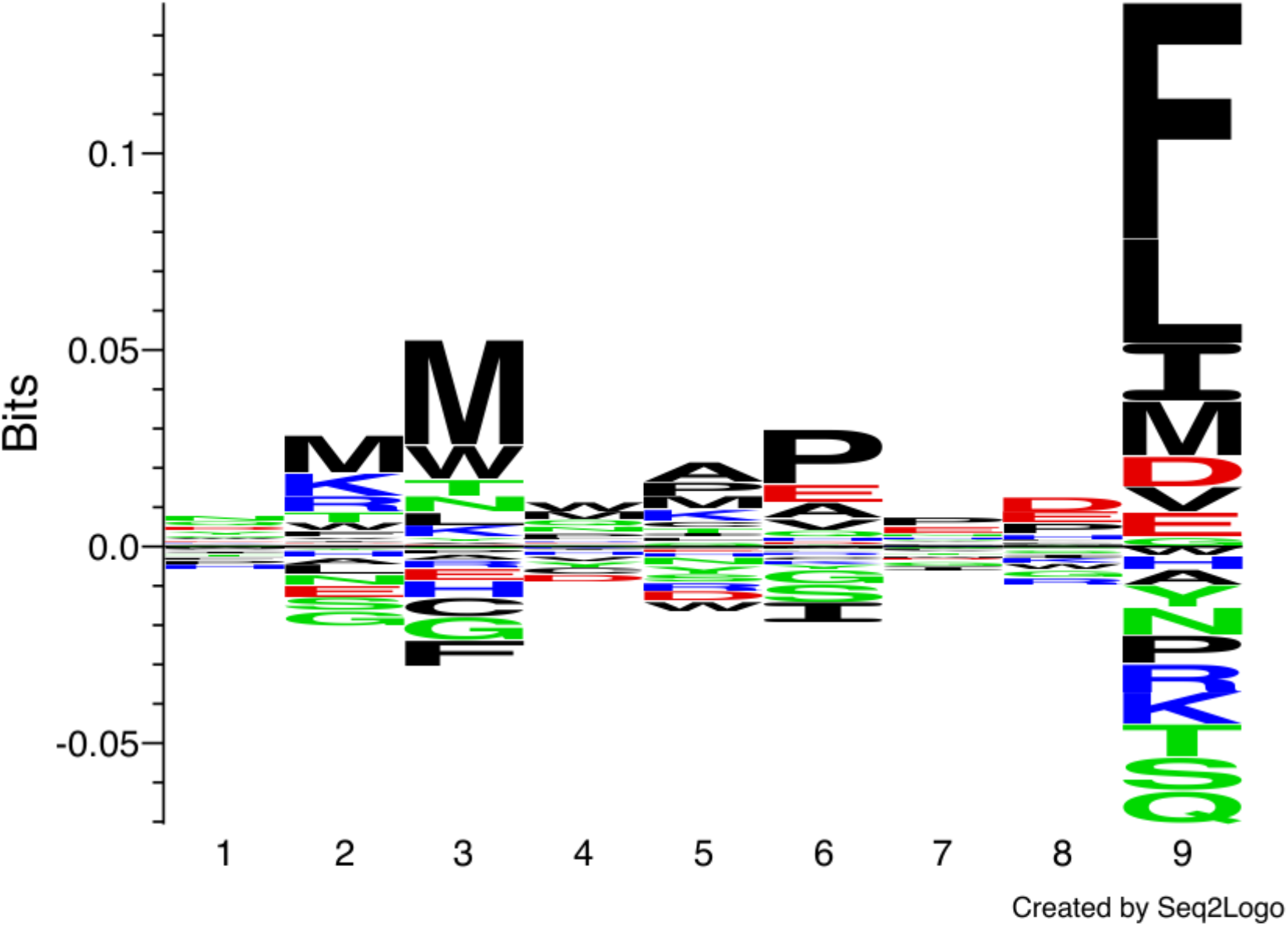
Peptide-binding logo for Acar3 based on PSCPL analysis using 9-mer peptide libraries. The peptide-binding logo was constructed with Seq2Logo 1.1 (Thomsen & Nielsen, 2012) based on the PSCPL-derived binding matrix ***(Figure S2)***. The relative height of each letter in the motif relates to the frequency of a given amino acid at that position in the bound 9-mer peptide.

### Peptide binding to Acar3 is governed by three footprints

To investigate the structural properties of Acar3, the Acar3-heavy chain (hc) and β_2_m were expressed in *Escherichia coli*, whereafter the Acar3-hc was refolded from inclusion bodies in the presence of soluble β_2_m and one of the three peptides (P1-P3), followed by crystallisation. High-quality crystal structures were obtained for P2 and P3 in space group P2_1_2_1_2_1_ at resolutions of 2.15 Å and 2.25 Å, respectively ***(Table 1, Figure S3)***. The solutions contained one peptide-Acar3 complex in the asymmetric unit, which consists of the Acar3-hc, β_2_m and either of the two peptides.

**Table 1:** Data collection and refinement statistics of Acar3·β2m in complex with peptide 2 and 3 (P2, P3).

| | [Acar3·β2m·P2] | [Acar3·β2m·P3] |
| --- | --- | --- |
| **Data collection** |  |  |
| Collection source | EMBL P14 | SLS X06SA |
| Space group | P2<sub>1</sub>2<sub>1</sub>2<sub>1</sub> | P2<sub>1</sub>2<sub>1</sub>2<sub>1</sub> |
| **Unit cell parameters** |  |  |
| a /b/c (Å) | 39.20/48.78/202.13 | 50.79/65.98/122.84 |
| $\alpha/\beta/\gamma$ (°) | 90.00/90.00/90.00 | 90.00/90.00/90.00 |
| Resolution (Å) <sup>a</sup> | 67.38 - 2.15<br>(2.21 - 2.15) | 46.94 - 2.25<br>(2.32 - 2.25) |
| Total reflections | 286,525 | 119,270 |
| Unique reflections | 22,044 | 20,176 |
| Completeness (%) <sup>a</sup> | 99.9 (99.8) | 99.5 (99.7) |
| Redundancy <sup>a</sup> | 13.0 (13.4) | 5.9 (5.7) |
| R<sub>merge</sub> (%) <sup>a</sup> | 12.8 (136.7) | 13.0 (111.4) |
| R<sub>pim</sub> (%) <sup>a</sup> | 5.2 (55.5) | 8.3 (74.9) |
| $\langle I/\sigma(I) \rangle$ <sup>a</sup> | 12.1 (1.9) | 8.1 (1.7) |
| CC(1/2) <sup>a</sup> | 0.999 (0.848) | 0.995 (0.418) |
| **Refinement** |  |  |
| R<sub>cryst</sub> (%) | 17.98 | 19.75 |
| R<sub>free</sub> (%) | 22.94 | 23.10 |
| Total number of atoms | 3338 | 3177 |
| **Average B-values (Å<sup>2</sup>) and (# of atoms)** |  |  |
| Acar3_hc | 53.6 (2,212) | 58.7 (2,184) |
| β2m | 51.9 (872) | 59.1 (841) |
| Peptide | 43.6 (75) | 62.8 (71) |
| Ligand/ion | 60.3 (10) | 81.7 (5) |
| Water | 50.7 (169) | 53.1 (76) |
| **r.m.s.d. from ideal** |  |  |
| Bond length (Å) | 0.0128 | 0.0134 |
| Bond angles (°) | 1.8366 | 1.9050 |
| **Ramachandran plot (MolProbity)** |  |  |
| Favored (%) | 96.88 | 96.85 |
| Allowed (%) | 2.86 | 2.89 |
| Outliers (%) | 0.26 | 0.26 |
| PDB code | 7ZQI | 7ZQJ |

The overall arrangement of the Acar3 complex resembles the commonly observed assembly of MHC-I complexes ***(Figure 2A)***. Six distinct pockets (A-F), previously identified in human HLAs [35], are present in the Acar3 peptide-binding groove (PBG), which is created by the α1 and α2 subunits with their α-helices lining the sides, and the shared β-sheet that forms the bottom of the crevice ***(Figure 2B, C; Figure S4)***.

**Figure 2:**
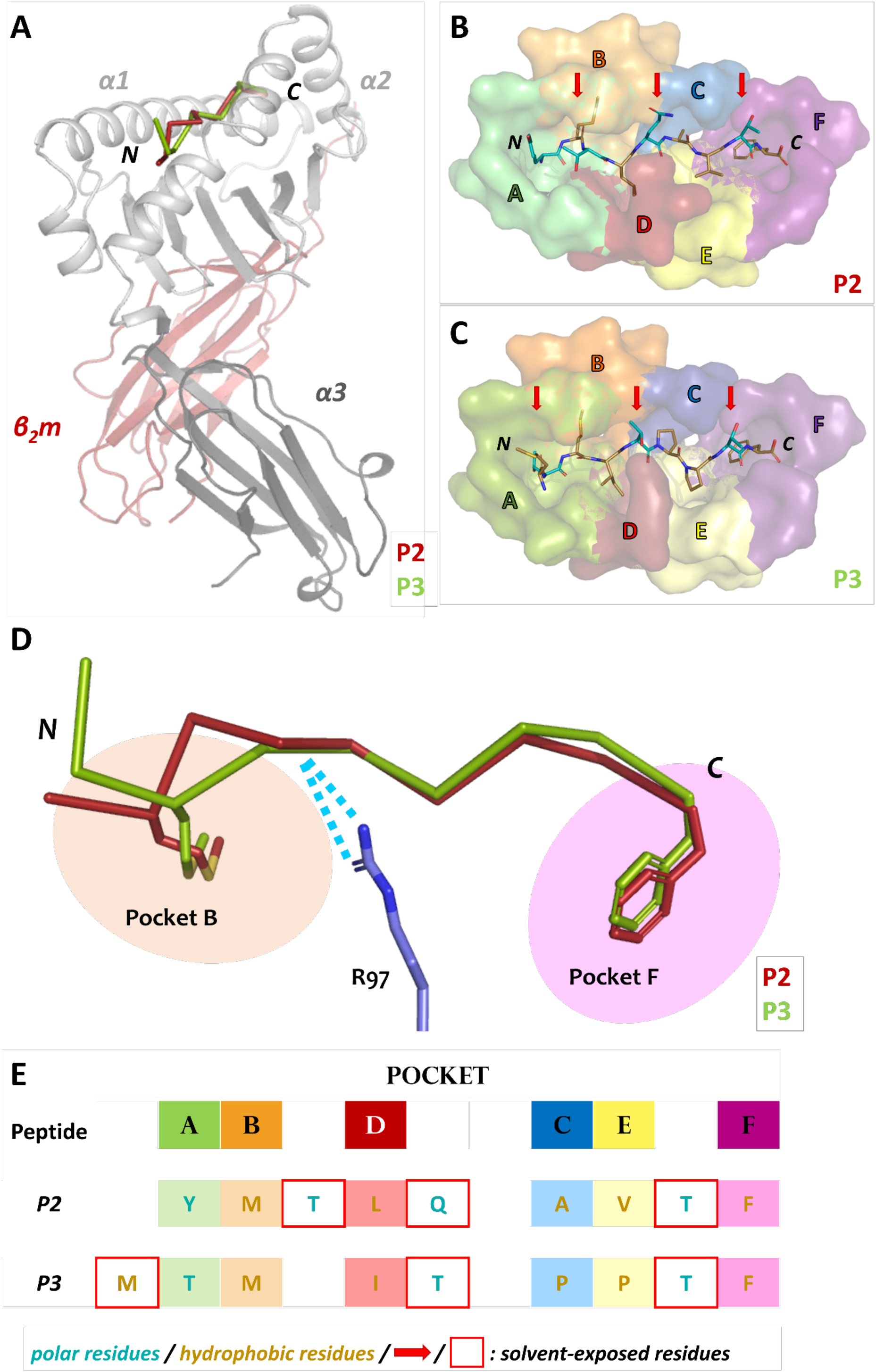
Overall architecture of the Acar3 MHC-I complex and pockets of the Acar3 peptide-binding groove (PBG) and peptide-binding mode. **(A)** Quaternary structure of the Acar3 complex with α1 and α2 represented in light grey, α3 in dark grey and β_2_m in red. The ribbon traces of the two peptides are shown in the PBG created by α1 and α2 (P2: red, P3: green). **(B-C)** Conformations of the peptides (shown as sticks) when bound to Acar3. Hydrophobic residues are shown in sand and polar residues are coloured teal. Solvent-exposed residues are indicated by red arrows. The pockets are shown as surfaces and coloured according to **Figure S5. (D)** Schematic representation of the three major anchor points within the Acar3 PBG. The peptides are shown as ribbons (P2: red, P3: green) and the side chains of the two anchor residues are represented as sticks. Arg97 from Acar3 is shown in stick representation in blue and interactions with the peptide backbones are indicated as blue dashes. **(E)** Schematic overview of buried (boxes coloured according to pockets) and solvent-exposed (indicated by a red frame) residues of the two peptides. The tabular arrangement illustrates the aa shifts of the peptides’ residues relative to each other upon binding to Acar3.

Comparing the binding modes of peptides P2 and P3 within the Acar3 PBG, it is evident that the five residues at positions 5-9 are well aligned in both structures, whereas their conformations differ substantially among their first four amino acids ***(Figure 2D, E)***. The Met anchor at position 2 of P2 is placed in pocket B of the Acar3 PBG. Simultaneously, the N-terminal Tyr_1_ residue is accommodated in the aromatic/hydrophobic environment created by the residues of pocket A ***(Figure)***. The N-terminus and the carbonyl oxygen of residue 1 establish hydrogen bonds with the three conserved Tyr residues (Tyr9, Tyr159, Tyr171) in pocket A ***(Table S2; Figure 3, top row)***. Conversely, the Met residue at position 3 (Met_3_) of P3 is positioned in pocket B which causes Thr at position 2 (Thr_2_) of the peptide to be accommodated in pocket A instead of the N-terminus. Consequently, the Thr_2_-side mimics the usual N-terminal interactions of antigenic peptides through engaging in a tight hydrogen-bond network with Tyr9 and Tyr171 ***(Table S3; Figure 3, top row)***. The N-terminus and side chain of Met_1_ stick out of the PBG to be solvent-exposed ***(Figure 2C, D, E)***. The side chain of Gln64 was found to play a prominent role in facilitating the unusual arrangement of P3 in the Acar3 PBG. While in the P2 structure Gln64 forms a hydrogen bond with the peptide-nitrogen of Met_2_, Gln64 creates a hydrogen bond with the carbonyl oxygen of M_1_ in the P3 complex ***(Figure 3, top row; Table S2, S3)***. The different orientation of the Gln64-side chain additionally decreases the negativity of the surface potential of pocket A, and thus contributes to the stabilisation of alternative N-terminal peptide conformations.

**Figure 3:**
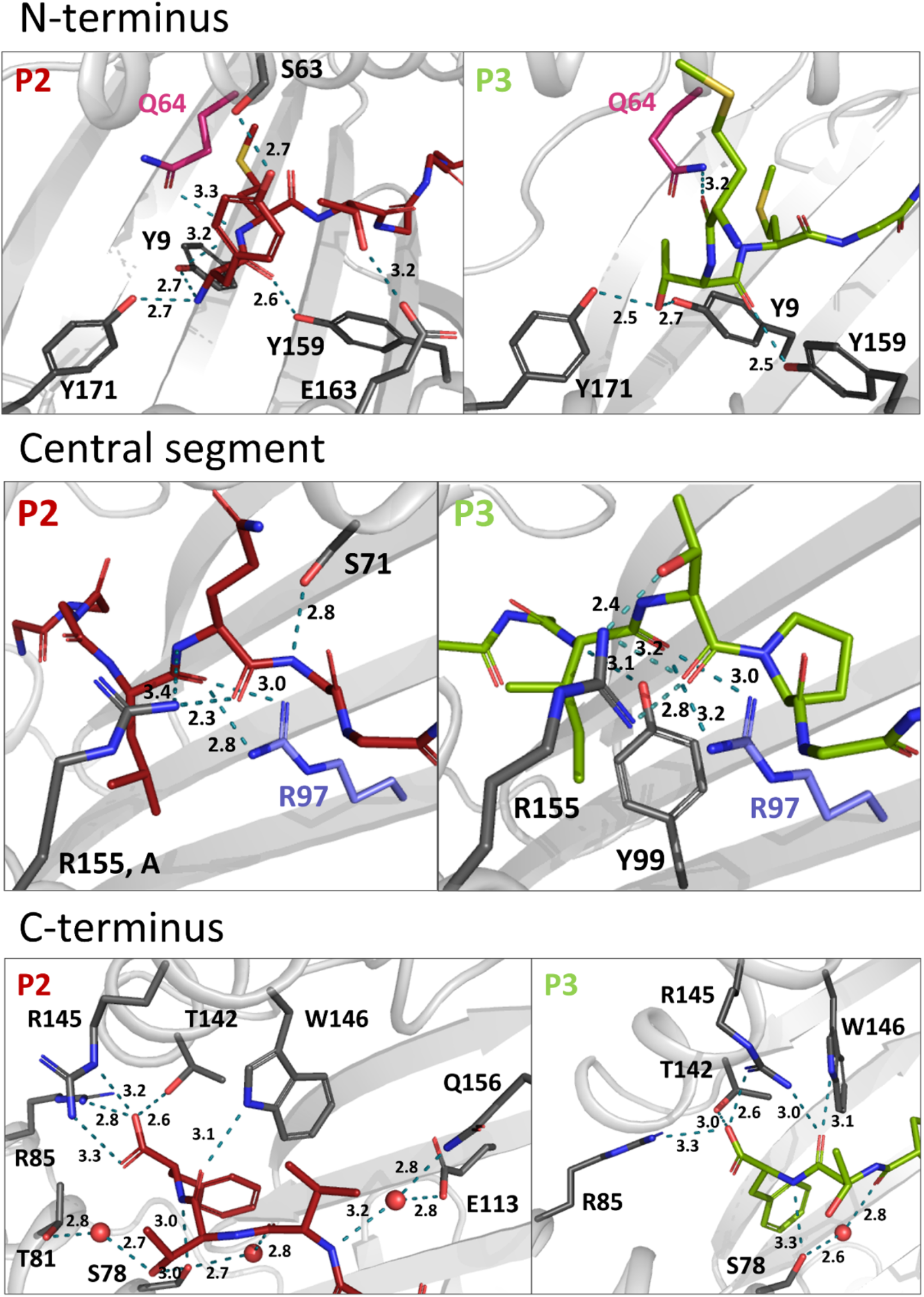
Hydrogen bonds of P2 and 3 within the Acar3 PBG. N-terminal **(top row)**, central **(middle row)** and C-terminal **(bottom row)** interactions are shown for P2 (red sticks) and P3 (green sticks). Residues that are involved in hydrogen bonds are shown in dark grey stick representation, and hydrogen bonds are indicated as teal dashed lines. Gln64 and Arg97 are highlighted in pink and blue, respectively. Water molecules are shown as red spheres and distances are given in Å. The side chains of peptide residues more distant from the relevant area have been omitted for clarity. A full list of interactions between P2 and 3 with Acar3 can be found in ***Tables S3-S4***.

In contrast to the elongated conformation of P3 resulting from the shifted position of the N-terminal Met-anchor residue, P2 adopts a bent conformation between the third and fifth residue ***(Figure 2D)*** with Thr_3_ being oriented towards the solvent, while Leu_4_ is inserted into pocked D ***(Figure 2B, E)***.

Following the conformational differences in the N-terminal segment, both peptides align starting from their residues at position 5. This convergence is facilitated through hydrogen-bonding interactions of the backbone-carbonyl oxygens of Leu_4_ in P2 and Ile_4_ in P3 with the guanidinium group of Arg97 which ascends from the bottom of the PBG ***(Figure 2D; Figure 3, middle row)***. Arg97 acts as a central, unspecific anchor of the peptide backbone and adopts an upright conformation, pointing directly towards the peptide. The orientation of Arg97 towards the centre of the PBG is greatly determined by the surrounding residues: In Acar3, the positioning of Arg97 is stabilised by interactions with its neighbouring residues Asp11 and Glu113 ***(Figure S5A)***. The combination of the three residues Asp11-Arg97-Glu113 forming a hydrogen-bond network does not occur in any HLA-I molecule. In HLA-B*15:01, which is closely related to Acar3 (discussed later) Arg97 is oriented towards the opposite direction and interacts primarily with Tyr74 (Tyr75 in Acar3), Asp114 (Glu113 in Acar3) and the C-terminal residue of the peptides that are presented by HLA-B*15:01 ***(Figure S5B)***. The proximity of Arg97 to pocket C in Acar3 may impact the preference for certain residues in the central part of the peptide as seen in the peptide-binding logo ***(Figure 1)***, such as the hydrophobic residues at position 6 of P2 (Ala_6_) and P3 (Pro_6_).

Residues Val_7_ (P2) and Pro_7_ (P3) are located in pocket E ***(Figure 2E)***, and the backbone-nitrogen atom Val_7_ participates in water-bridged hydrogen bonds with Glu113/Gln156 ***(Figure 3, bottom row)***. In both Acar3-peptide complexes, the aromatic side chain of Phe_9_ is deeply buried in pocket F with the backbone-NH and the carboxy terminus oriented towards the opening of the PBG. The Phe_9_-side chain participates in hydrophobic contacts with the pocket-F residues as well as in π-stacking interactions with Phe115 and Phe122. The C-terminal carboxy group of P2 establishes strong hydrogen bonds with Arg85 and Thr142, and slightly weaker hydrogen bonds with Arg145. Moreover, the backbone-NH of Phe_9_ (P2) forms a hydrogen bond with Ser78. P3 forms similar hydrogen bonds to Acar3 as observed for P2, except for the interactions with Arg145 and Thr81 ***(Figure 3, bottom row; Table S2, S3)***.

Overall, the peptide binding signature in Acar3 is determined by three structural features: 1) the adaptability of pocket A and the spaciousness of pocket B (discussed below) to permit great flexibility of the N-terminal part of the peptides, 2) the central backbone anchor Arg97 that imposes conformational uniformity on the C-terminal backbone trace of bound antigens, and 3) the narrow and specific pocket F, which allows tight binding of the C-terminal anchor residue of the peptides ***(Figure 2D)***.

### A unique composition of pocket B residues permits great N-terminal flexibility of peptides bound to Acar3

A search for P1, P2 and P3 sequences in the Immune Epitope Database (IEDB) revealed that in previous studies P1 has been found to bind HLA-B*15:01 [36, 37], while P2 shows affinity to a broader range of HLA-I molecules, such as HLA-A*24:02, HLA-A*24:03, HLA-A*69:01, HLA-B*15:01, HLA-B*27:05 and HLA-B*35:01 [36-38]. Studies of P3-binding to different HLA-I molecules revealed an even broader spectrum of binding partners, *i*.*e*. HLA-A*24:02, HLA-A*32:07, HLA-A*32:15, HLA-A*68:23, HLA-B*15:42, HLA-B*35:01, HLA-B*45:06, HLA-B*58:01, HLA-B*83:01 and HLA-C*04:01 [36, 37, 39]. To evaluate which of the HLA molecules that bind P1, P2 and P3 obtained from the IEDB are most similar to Acar3, we prepared a distance tree based on peptide-binding motifs using *in-silico* MHC-binding predictions [40] ***(Figure 4A)***. The five most closely related HLA-I molecules to Acar3, which bind at least one of the three peptides, are HLA-A*32:07, HLA-A*32:15, HLA-A*24:02, HLA-A*24:03 and HLA-B*15:01. Peptide-binding logos from more distant HLA molecules (available on the *NetMHCpan 4*.*0 Motif Viewer* website) which were found to bind P2 and/or P3, showed either ambiguous or inconclusive residue preferences at position 2 or a preference for Thr, which occupies the second position of P3 and the third position of P2. An alignment of the residues (including key residues) that flank pocket B [41] in Acar3 and the five HLA-I molecules reveals a unique amino-acid signature of this pocket for Acar3 ***(Figure 4B)***. The key residues define the shape and depth of the pockets and establish interactions with the peptides’ anchor residues. Those HLA-I molecules most closely related to Acar3, whose structures are available in the PDB, *i*.*e*. HLA-A*24:02 and HLA-B*15:01, reveal a rather narrow and elongated shape of pocket F similar to Acar3, thereby permitting little conformational flexibility of the C-terminus. In contrast, compared to HLA-A*24:02 and HLA-B*15:01, pocket B of Acar3 is wider and more open towards the PBG, which is the result of small residues at positions 11 (Asp) and 71 (Ser), segregating pocket B from pocket C ***(Figure 4C, top row)***. Moreover, with respect to all known classical and non-classical HLA class-I molecules, a Gly residue at position 68 (67 in HLA) is a unique feature of Acar3 and has particular impact on the depth and width of pocket B. In Acar3, the peptide’s anchor residue accommodated in pocket B has thus greater conformational freedom which results in an increased flexibility of the N-terminal half of the peptide. In addition to Met as the preferred anchor residue, pocket B provides sufficient space to accommodate residues of secondary importance, *e*.*g*. Trp, as occurring in the peptide-binding logo ***(Figure 1)***. In contrast, the anchor residues located in the B pockets of HLA-A*24:02 and HLA-B*15:01 show less positional deviation among different structures – even when anchor residues of different chemical properties and peptides of different lengths are considered ***(Figure 4C, bottom row)***. This is reflected by the root-mean-square deviation (r.m.s.d.) values among the N-terminal regions (aa 1-3) of the peptides bound to Acar3 being twice as high as in HLA-A*24:02 and HLA-B*15:01-peptide complexes. In contrast, the relative flexibility among the central amino acids (aa 4-6) of peptides bound to HLA-A*24:02 and HLA-B*15:01 is significantly higher (≈ 2-fold) than among Acar3-bound peptides ***(Table S4)***.

**Figure 4:**
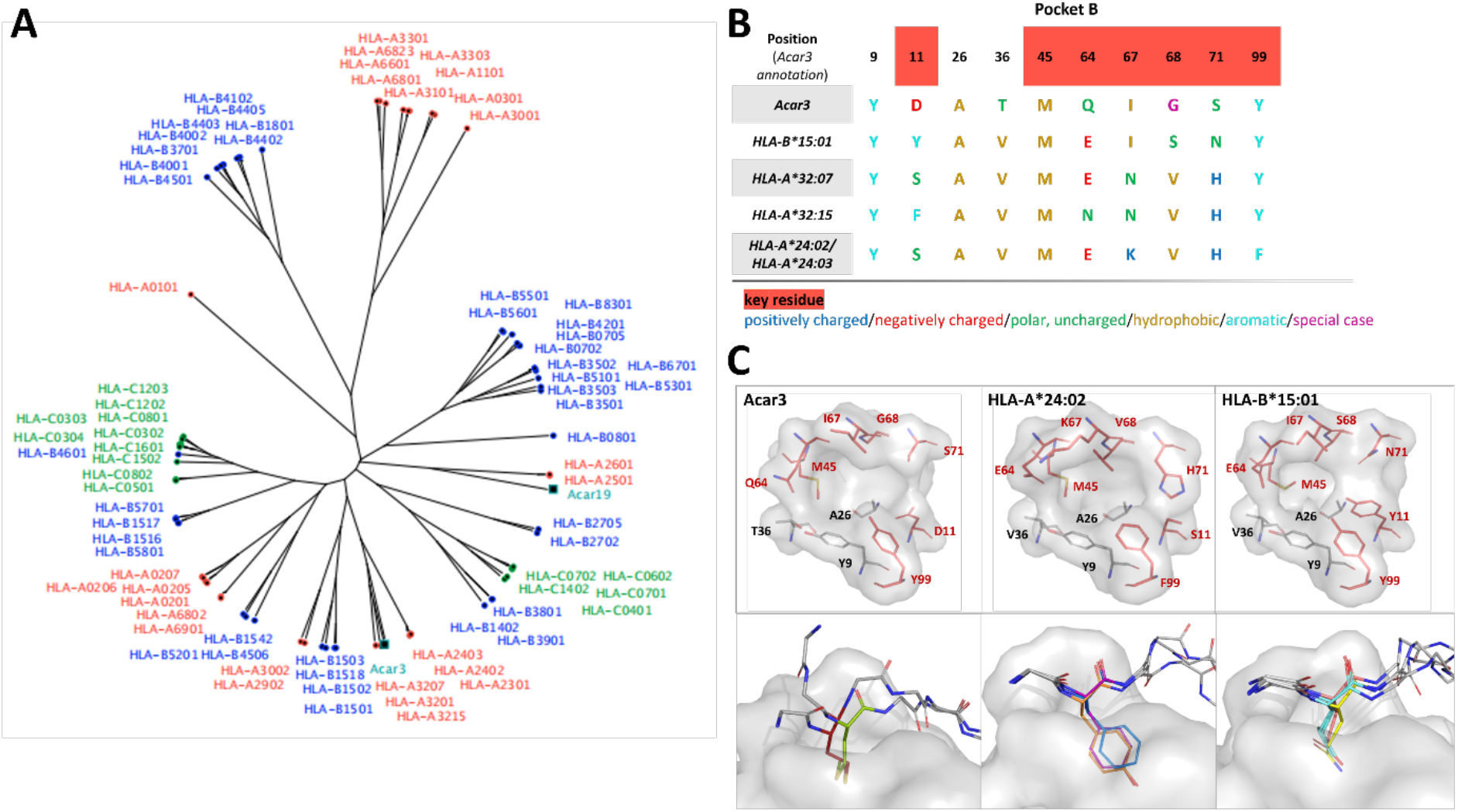
Comparison of Acar3 with phylogenetically close HLA-I molecules. **(A)** Phylogenetic tree including Acar3, Acar19 and HLA class-I molecules. **(B)** Comparison of pocket B residues of Acar3 with its closest HLA-I molecules. Key-residue positions are indicated in red. **(C, top row)** The pockets from **(B)** shown as surface (grey) and stick representation with key residues coloured red. **(C, bottom row)** Positional variance of anchor residues accommodated in pocket B of Acar3, HLA-A*24:02 and HLA-B*15:01. The pocket is shown as grey surface, the peptides are shown as sticks with the anchor residue in colour and the surrounding residues shown in grey and without side chains. For Acar3, the anchor residues are depicted in red (M_2_ (P2)) and green (M_3_ (P3)). From the structures of HLA-A*24:02 and HLA-B*15:01 available in the PDB, three representatives have been chosen each. HLA-A*24:02: 2BCK, chain A, B and C (Y_2_ in orange), 3I6L (F_2_ in blue) and 4F7M chain A, B and C (Y_2_ in magenta); HLA-B*15:01: 1XR8 (E_2_ in salmon), 1XR9 (L_2_ in cyan) and 5VZ5 (Q_2_ in yellow).

## Discussion

Great reed warblers are long-distance migratory songbirds that successfully handle pathogens at breeding, stop-over and wintering sites [42]. It has been suggested that their highly duplicated MHC-I genes are associated with particularly well-developed immune adaptations [43]. Our work presents the first structure of an MHC-I molecule from a wild songbird in complex with three different antigenic peptides and elucidates the structural basis of a unique peptide-binding mode. Although the overall architecture of the Acar3 peptide-binding groove comprising six distinct pockets closely resembles the characteristics of HLA class-I molecules, Acar3 does not completely resemble any supertypes defined for HLA-I molecules [41]. Together with the unique amino-acid composition and shape of pocket B, the central Arg97 residue strongly contributes to defining the Acar3-peptide binding mode. Whereas Arg97 draws the central part of the bound peptides closer to the bottom of the PBG, thereby contrasting the conformation usually observed in peptide-HLA-I complexes, pocket B enables an outstanding N-terminal flexibility with Gln64 permitting a shifted arrangement of the peptides’ N-terminal part. This observation puts the necessity of a Glu residue at position 63 in porcine MHC I (SLA-1*0401) and in HLA class I (corresponding to Gln64 in Acar3) for the N-terminal extension presentation mode as proposed by a recent study [44] into perspective, and suggests that N-terminally extended peptide binding can be facilitated in different ways across vertebrate species. With respect to the possibility for either Met_2_ or Met_3_ to be accommodated in pocket B of Acar3, it is likely that positions 2 and 3 in the peptide-binding logo ***(Figure 1)*** represent a combined anchor rather than two distinct positional amino-acid specificities. In addition to the preference for Met at both positions, pocket B provides sufficient space to accommodate a Trp residue. The slightly negative surface potential of the cavity could furthermore permit the insertion of Lys and Arg, which represent subordinate anchor residues at position 2 of the peptide-binding logo. The increased residue preference at position 6 of the peptide-binding logo possibly results from the role of Arg97, which anchors the bound peptides to the centre of the PBG. However, the different properties of the amino acids at position 6, *i*.*e*. Pro, Glu and Ala, probably reflect the unspecific nature of this interaction through the formation of hydrogen bonds with the peptide backbone.

The preferred anchor residues assigned to pocket B, Met and – to a lesser extent – Trp, usually occur at low frequency in the proteomes of living organisms, including potential pathogens [45-50] ***(Figure S6)***. Those residues are among the amino acids with the highest metabolic synthesis costs [45, 46, 51], which potentially causes them to be present almost exclusively at specific sites within proteins. Many different criteria have been established to predict the mutational probabilities of amino-acid residues in proteins, including protein stability, genetic code and physicochemical similarities of amino acids [52-54]. Due to their distinct properties with respect to *e*.*g*. resilience to oxidative stress (Met) and protein folding (Trp) [45, 55-58], a substitution of those residues within the context of a protein will likely result in less stable – probably even less functional – variants, and thus impose a fitness cost on the pathogen. Therefore, the mutation rate of Met and Trp is comparably low [52, 59], impeding substitutions of potential anchor residues that would prohibit binding of pathogen-derived peptides to Acar3. The conformational variability in the N-terminal halves of Acar3-binding peptides results in two potential positions (2 or 3) of the anchor residue within the peptide sequence. This increases the repertoire of antigens harbouring rare amino acids like Met or Trp that can be presented by Acar3. However, the accommodation of an anchor residue such as Met at position 3 of the peptide in pocket B will probably depend on the presence of a small, polar residue, *e*.*g*. Thr or Ser, at position 2 acting as a mimic for the N-terminal amino group.

Two recent studies reported on an N-terminally shifted binding mode of the HIV Gag epitope TW10 (TSTLQEQIGW) to HLA-B*57:01 and HLA-B*58:01 [60, 61], closely resembling the non-canonical binding of P3 to Acar3 presented in this work. In both studies, the N-terminal residue of the TW10 peptide is solvent exposed while Ser at position two occupies the actual position of the amino group in pocket A. On the contrary, an HIV-escape mutant, TW10-T3N (TS**N**LQEQIGW), leads to a canonical binding mode of this peptide variant with Thr at position one being accommodated in pocket A of HLA-B*57:01. The N-terminal register shift causes major conformational changes in the C-terminal part of the bound peptide, thereby attenuating antigen recognition of the escape mutant by the NK receptor KIR3DL1 [60]. Intriguingly, in Acar3 no such C-terminal conformational changes caused by the N-terminal register shift can be observed, which emphasises the particular role of the central Arg97 acting as an unspecific anchor for the peptide backbone. The non-distinctive nature of this central anchoring point with respect to peptide sequence renders the antigen presenting system less susceptible towards escape mutations in pathogens. Hence, the unique peptide-binding mode of Acar3 could have important implications for maintaining the antigen presentation and for hindering immune evasion by pathogens: Both, the expansion of the peptide repertoire through permitting N-terminal flexibility, as well as putative pathogen escape mechanisms may not impair antigen recognition by NK receptors that interact with the C-terminal part of the presented peptide. These include receptors with similar functions to killer immunoglobulin-like receptors (KIR) which have been identified in many vertebrate species [62-65], and are thus likely to be present also in songbirds.

Although the majority of peptides presented by MHC molecules are of self-origin, this is unlikely in the case of the three top-binding peptides to Acar3, since they were not found within the great reed warbler exome. It is therefore more likely that they stem from either viral or bacterial pathogens capable of invading cells, which explains their presentation on MHC class-I molecules rather than MHC-II molecules. Using the example of the role of HLA-B*57 in HIV, it has been demonstrated that the interplay between the innate immune response mediated by KIR3DL1 recognition and the adaptive immunity through the activation of CD8^+^ T cells enhances the protection against an unfavourable disease outcome [60, 66]. Acar3 may thus have a role at the interface of innate and adaptive immunity and could be an example of a sophisticated mechanism in migratory birds to counter immune evasion of different pathogens.

## Materials and methods

### Amino acid sequences of Acar3, Acar5, Acar19 and Beta-2-microglobulin

The Acar3, Acar5 and Acar19 heavy-chain (hc) sequences have been published previously [31, 67]. To amplify the *β*_***2***_*m* gene, degenerated PCR-primers were designed from transcriptomic *β*_***2***_*m* sequences in willow warbler (*Phylloscopus trochilus*, NCBI SRA Accession no. SRA056327 [68], house sparrow (*Pado*, NCBI SRA Accession no. SRP012188, [69]) and zebra finch (*T. guttata*, GenBank accession no; DQ215661 and DQ215662, [70]). A 533-bp long *β*_*2*_*m* fragment spanning the whole mature peptide region and partially the 3’ UTR region was amplified with the degenerated forward primer P162 5’ CGGGGWGCCCTGGCGCTC 3’ and the reverse primer P149 5’ GGCCTGCAGACCTCCCTTGA 3’ using *Acar* cDNA template from two great reed warbler individuals. Each PCR reaction contained 2 μl diluted cDNA (for details on RNA extraction and cDNA preparation [71]), 0.2 μM of each primer, 1.5 mM MgCl_2_, 1.25 U AmpliTaq DNA Polymerase and 1x GeneAmp Buffer II (Applied Biosystems) in a total volume of 25 μl. The following cycling parameters were used: 94 °C for 3 min and then 35 cycles of 94 °C for 30 s, 58 °C for 30 s and 72 °C for 90 s. A final extension step at 72 °C for 15 min was applied. PCR products were Sanger sequenced in the forward and reverse direction using standard procedures (BigDye terminator Cycle Sequencing kit v.3.1, Life Technologies) on an ABI PRISM 3130 genetic analyser (Applied Biosystems). The resulting sequences were inspected and manually edited with the software Geneious (v.5.5, Biomatters). A 495 bp-long partial *Acar β*_***2***_*m* transcript was successfully sequenced from great reed warbler cDNA (*Acar β*_*2*_*m*, GenBank Accession no. KM096440). No polymorphic sites were detected within or between the two sequenced individuals. The *Acar-β*_***2***_*m* fragment included partial signal peptide information (39 bp), the whole predicted mature peptide (297 bp) and partial 3’ UTR (156 bp).

### Protein production (heavy chain (hc) and β_2_m) for Scintillation Proximity Assays (SPA)

*Acar* MHC-I heavy chains (residues 1-272) containing a C-terminal histidine-affinity tag for purification and biotinylation substrate peptide (BSP) for biotinylation were produced recombinantly and purified as previously described [72]. In brief, the proteins were overexpressed in *Escherichia coli*, resulting in the formation of inclusion bodies. After mechanical cell lysis, inclusion bodies were isolated by centrifugation and dissolved in 8 M urea, 25 mM Tris/HCl, pH 8.0, and purified by immobilised metal (Ni^2+^) affinity chromatography (IMAC), hydrophobic interaction chromatography and size exclusion chromatography. *In-vivo* biotinylation of the BSP-tagged MHC-I hc was achieved by co-expression of the BirA enzyme.

*Acar* β_2_m containing an N-terminal histidine-affinity tag and a factor Xa cleavage site, was produced by overexpression in *Escherichia coli* and the inclusion bodies were isolated and dissolved in 8 M urea as described above. The protein was first purified by IMAC in 8 M urea, 25 mM Tris/HCl, pH 8.0. The purified and denatured β_2_m was folded by drop-wise dilution into a non-denaturing buffer (300 mM urea, 25 mM Tris/HCl, pH 8.0) over 24 hours. The folded β_2_m protein was then treated with Factor Xa to remove the N-terminal affinity tag. Thereafter, cleaved β_2_m was separated from the uncleaved fraction and the affinity tag by IMAC, and lastly purified by size-exclusion chromatography (SDX200-PG).

### Generation of a 9-mer Positional Scanning Combinatorial Peptide Library (PSCPL)

Peptides of the 9-mer PSCPL were synthesised with an equimolar mixture of 19 of the 20 amino acids found in proteins (excluding Cys) that are randomly coupled in eight positions, with one specific amino acid (including Cys) being fixed at one position. This yields nine libraries for each amino acid, one library for each position of the peptide. 20 amino acids thus give a total of 20 × 9 = 180 peptide libraries. In one synthesis the amino acid pool was used in all nine positions, giving a total of 181 individual peptide libraries, for further details see [34, 73, 74].

### Scintillation Proximity Assay (SPA) based peptide-MHC-I dissociation assays

Both to determine the peptide length preference of two different MHC-I molecules, Acar19 and Acar5, and the binding motif of Acar19, Acar5 and Acar3, peptide-Acar (pMHC-I) dissociation was measured using a scintillation proximity assay (SPA) as described in [34, 74]. To determine the preferred length, we evaluated peptide binding in terms of the amounts of complexes formed with peptides of different lengths in an unbiased way, and thus employed peptide libraries ranging from 7 to 13 residues in length. Since the motif characterisation in this study only focussed on Acar3, Acar3-complex stability was determined for nonameric peptides from the 9-mer PSCPL using a scintillation proximity assay (SPA) [72, 74]. Briefly, denatured and biotinylated Acar3-hc was diluted into PBS/0.1 % Lutrol® F68 containing 10μM of peptide and trace amounts of ^125^I radiolabelled β_2_m in streptavidin coated scintillation microplates (FlashPlate®, Perkin Elmer, SMP103001PK or SMP410001PK). Plates were incubated over night at 18 °C to attain complex folding. Dissociation was initiated at time point zero (Y_0_) by addition of unlabelled β_2_m and scintillation was measured continuously for up to 24 hours at 37 °C in a TopCount NXT liquid scintillation counter (Packard). The resulting dissociation data for each peptide sub-library was used to calculate a relative binding value (RB-value) by dividing the approximated area under the curve (AUC) of the sub-library with the AUC of the reference library. A matrix using the RB-values was then generated where an RB-value ≥ 2 defines a favoured amino acid at a specific position, and an RB-value of ≤ 0.5 defines a disfavoured amino acid at a specific position [73, 74]. Thus, the matrix defines the peptide-binding properties of the Acar3 molecule. The logo of the PSCPL-derived Acar3-binding motif was made using the Seq2Logo 1.1 server as P-Weighted Kullback-Leibler logos [75].

Using the matrix obtained from the PSCPL, a predicted Acar3-binding score can be calculated for a given 9-mer peptide sequence by multiplying the RB values for each position of a specific peptide, *e*.*g*. (aa Pos. 1 RB)*(aa Pos. 2 RB)*(aa Pos. 3 RB)*(aa Pos. 4 RB)*(aa Pos. 5 RB)*(aa Pos. 6 RB)*(aa Pos. 7 RB)*(aa Pos. 8 RB)*(aa Pos. 9 RB) [73]. The rank score was calculated for some 9500 *in-house* 9-mer peptides and the top 94 scoring peptides were selected for binding studies using SPA as described above.

The three peptides that yielded pMHC-I complexes with the highest stability in the SPA were peptide 1 (P1, AMSAQAAAF), peptide 2 (P2, YMTLQAVTF) and peptide 3 (P3, MTMITPPTF) ***(Table S1)***.

### Functional clustering

The *Acar* MHC-I and human HLA-I allomorphs were clustered using the *MHCcluster* method [76], which predicts and functionally clusters the peptide-binding specificities of the MHC-I allomorphs. The *MHCcluster* method used here was based on the retrained version of the *NetMHCpan* method [77], trained including a small set of peptides with measured half-lives for Acar3 (188 peptides) and Acar19 (275 peptides) complexes ***(Table S5)***. *MHCcluster* estimates the functional similarities between any two MHC-I allomorphs in the analysis by correlating the union of the predicted top 10 % of strongest binding peptides for each of the defined allomorphs. If two MHC-I allomorphs were predicted to have a perfect overlap regarding their peptide-binding specificities, the similarity was defined as 1, and if there was no overlap at all, the similarity was defined as 0. The distance matrix was converted to a distance tree using UPGMA clustering. To estimate the significance of the MHC-I distance tree, 1000 distance trees were generated using the bootstrap method. The bootstrapping was performed at the peptide level, *i*.*e*. for each distance tree a new set of 100.000 peptides and the correlating prediction was selected from the original pool of 100.000 peptides with replacements. The trees were then summarised, and a consensus tree was made with branch bootstrap values.

### Protein production and refolding for structural studies

The Acar3-hc (residues 1-272) was cloned into a pET26b(+) vector, and electrocompetent *E. coli* TUNER (DE3) cells were transformed with the plasmid. Subsequent protein expression was performed using BD Difco LB broth Miller supplemented with 50 μg/ml kanamycin. Cells were grown at 37 °C, 120 rpm, and when an OD_600_ of 0.8 was achieved, IPTG was added to a final concentration of 1 mM. Four hours after induction, cells were harvested, and the pellets were stored at -80°C. The inclusion bodies containing the Acar3-hc, were isolated from the cells and washed according to previously published protocols [78, 79]. Finally, the inclusion bodies were solubilised in 20 mM Tris/HCl, pH 8.0, 8 M urea, 1 mM EDTA, 10 mM DTT (hereafter referred to as “IB buffer”) and stored at -80 °C until further use.

A synthetic *β*_*2*_*m* gene that was codon-optimised for *E. coli* was ordered from GenScript and cloned into the pET-26b(+) vector. The gene was then moved to the expression vector pNIC28-Bsa4 using ligase independent cloning (LIC). The resulting pNIC28-b2m construct encodes β_2_m with an N-terminal (His)_6_-tag that can be removed by TEV cleavage, leaving a single N-terminal Ser on β_2_m. Electrocompetent *E. coli* TUNER (DE3) cells were transformed with the plasmid and expression was carried out in TB medium supplemented with 50 μg/ml kanamycin. The culture was started at 30 °C, 120 rpm, and at an OD_600_ of 0.5, the temperature was lowered to 18 °C. At OD_600_ = 0.9, IPTG was added to a final concentration of 0.1 mM. The cells were harvested 20 hours after induction and the pellets were solubilised in 50 mM Tris/HCl, pH 8.0, 300 mM NaCl, 20 mM imidazole. Cells were disrupted mechanically through sonication or application of pressure and the lysate was applied to a Ni^2+-^affinity column using a gradient of 20-500 mM imidazole. Afterwards, the protein was purified by size-exclusion chromatography (SEC) in 20 mM Tris/HCl, pH 8.0, 150 mM NaCl, 1 mM EDTA, concentrated and stored at -80 °C until further use.

Purified β_2_m was added to 200 ml of 100 mM Tris/HCl, pH 8.0, 400 mM L-Arg, 1 mM EDTA, 5 mM GSH, 0.5 mM GSSG, 0.5 mM PMSF at 4 °C under gentle stirring to a final concentration of 2 μM. Denatured Acar3 (solubilised in IB buffer) was premixed with P2 (solubilised in DMF) or P3 (solubilised in 50 % (v/v) water + 50 % (v/v) IB buffer) in a concentration that would result in final concentrations of 1 μM Acar3 and 10 μM peptide upon addition to the β_2_m-containing refolding buffer. The peptides used for the refolding experiments were purchased from Peptides International Inc. with a purity level of ≥ 95 %. Refolding was initiated by the dropwise addition of 1/3 of the volume of the Acar3-peptide mixture to the β_2_m-containing refolding buffer under continuous stirring. After 12 and 24 h each, again 1/3 of the Acar3-peptide mixture were added and stirring continued. After 48 h of incubation, the refolding mixture was transferred to 4 l of 50 mM Tris/HCl, pH 8.0, 150 mM NaCl, 0.5 mM EDTA and dialysed for 2 h. Then, the mixture was moved to 4 l of 50 mM Tris/HCl, pH 7.5, 150 mM NaCl and dialysed for approx. 10 h. The dialysate was concentrated and applied to a HiLoad 16/600 Superdex 200 column using 20 mM Tris/HCl, pH 7.5, 150 mM NaCl (SEC buffer) and the fractions containing the complex were pooled and concentrated.

### Crystallisation and structure determination of [Acar3·β_2_m] in complex with P2 and P3

Prior to crystallisation, the refolded complexes were diluted in SEC buffer to final concentrations of 5 mg·ml^−1^ ([Acar3·β_2_m·P2]) or 10 mg·ml^−1^ ([Acar3·β_2_m·P3]) and centrifuged for 10 min at 16100 x g. Crystallisation was performed at 293 K, using the hanging-drop vapour-diffusion method at a protein-to-reservoir ratio of 1:1 in the drop. To obtain crystals of sufficient size and quality, the [Acar3·β_2_m·P2] crystallisation drops were subjected to streak seeding, using a seeding solution from previously obtained crystals that were smashed by sonication. Crystals of [Acar3·β_2_m·P2] grew in 100 mM Hepes/MOPS, pH 7.0, 60 mM divalent mix (20 mM MgCl_2_/40 mM CaCl_2_) and 32.5 % precipitant mix (25 % (v/v) MPD/25 % (w/v) PEG 1000/25 % (w/v) PEG 3350) and were flash-frozen in N_2_ (l) without cryo-protection. Crystals of [Acar3·β_2_m·P3] were obtained in 100 mM MES, pH 6.0, 6 % (v/v) Tacsimate, pH 6.0 and 25 % (w/v) PEG 4000, cryo-protected with 35 % (v/v) ethylene glycol and flash-frozen in N_2_ (l).

Diffraction data were collected at DESY (PETRA III), EMBL Hamburg, Germany, at beamline P14 ([Acar3·β_2_m·P2]), and at SLS, PSI Villigen, Switzerland, at beamline X06SA ([Acar3·β_2_m·P3]). The data were processed with XDS [80] and further steps were carried out using programs from the CCP4 program suite [81]. Molecular replacement was performed with PHASER [82] using an MHC-I molecule from duck (*Anas platyrhynchos*, PDB: 5GJX) as search model for the [Acar3·β_2_m·P3] structure. Thereafter, for [Acar3·β_2_m·P2], the previously solved structure of [Acar3·β_2_m] was used as search model. Structure refinement was carried using REFMAC [83] and phenix.refine [84], and model building was performed with Coot [85, 86].

The protein structures presented in this article has been submitted to the Protein Data Bank (http://www.rcsb.org/pdb/home/home.do) under accession numbers 7ZQI and 7ZQJ.

## Acknowledgements

This study was funded by the European Research Council (ERC) under the European Union’s Horizon 2020 research and innovation programme grant 679799 to HW. We thank Prof. Dr. Caroline Kisker for critical reading of the manuscript and valuable suggestions; and we are grateful to Prof. Dr. Caroline Kisker and Prof. Dr. Hermann Schindelin for helpful discussion of the structural data. Moreover, we thank Dr Michael Rasmussen, Dr Maria Strandh and Dr Elna Follin for help with labwork, analyses and the scientific discussions. We thank the staff of the beamline P14 at Petra III at the Deutsches Elektronen-Synchrotron (DESY) in Hamburg, and at beamline X06SA at Swiss Light Source (SLS) in Villigen for excellent support. Moreover, we thank HW’s horse Maggan for providing resources (hair) for streak seeding.

## Author contributions

Conceptualization: HW, KL-P and KP conceived the study. Methodology: SB, MN, KL-P, and HW provided methodologies in their respective expertise. Formal analysis: SE, SM, SB, and MN analyzed data related to their respective expertise. Investigation:SE and SB performed the experiments and collected the data. Resources: HW and KL-P provided study material, reagents and materials. Writing: SE provided the first draft of the manuscript and all co-authors contributed with reading, revising and commenting on the manuscript until the final version. Visualization: SE, SM, SB, and MN prepared the figures and tables. Supervision: HW took the main leadership responsibility for the research activity. Project administration: HW coordinated the responsibility for the research activity planning and execution. Funding acquisition: HW was responsible for the financial support for the project leading to this publication.

## References

1. Jetz, W., et al., The global diversity of birds in space and time. Nature, 2012. 491(7424): p. 444–8.

2. Pigot, A.L., et al., Macroevolutionary convergence connects morphological form to ecological function in birds. Nat Ecol Evol, 2020. 4(2): p. 230–239.

3. Pigot, A.L., C.H. Trisos, and J.A. Tobias, Functional traits reveal the expansion and packing of ecological niche space underlying an elevational diversity gradient in passerine birds. Proceedings of the Royal Society B: Biological Sciences, 2016. 283(1822): p. 20152013.

4. del Hoyo, J., et al., Handbook of the Birds of the World Alive. 2016, Lynx Edicions: Barcelona.

5. Clark, N.J., S.M. Clegg, and M. Klaassen, Migration strategy and pathogen risk: non-breeding distribution drives malaria prevalence in migratory waders. Oikos, 2016. 125(9): p. 1358–1368.

6. Mendes, L., et al., Disease-limited distributions? Contrasts in the prevalence of avian malaria in shorebird species using marine and freshwater habitats. Oikos, 2005. 109(2): p. 396–404.

7. Merino, S., et al., Haematozoa in forest birds from southern Chile: Latitudinal gradients in prevalence and parasite lineage richness. Austral Ecology, 2008. 33: p. 329–340.

8. Williams, C.W., Evolution in Health and Disease, second edition Stephen C. Stearns, Jacob C. Koella (Eds). Oxford University Press Inc., New York, 2008. 400 pp., paperback, ISBN: 978-0-19-920746-6 (US$69.95). Transactions of The Royal Society of Tropical Medicine and Hygiene, 2008. 102(11): p. 1168–1168.

9. Bordes, F., J.F. Guégan, and S. Morand, Microparasite species richness in rodents is higher at lower latitudes and is associated with reduced litter size. Oikos, 2011. 120(12): p. 1889–1896.

10. Nunn, C.L., et al., Latitudinal gradients of parasite species richness in primates. Diversity and distributions, 2005. 11(3): p. 249–256.

11. Goldstein, T., et al., The discovery of Bombali virus adds further support for bats as hosts of ebolaviruses. Nat Microbiol, 2018. 3(10): p. 1084–1089.

12. Calisher, C.H., et al., Bats: important reservoir hosts of emerging viruses. Clin Microbiol Rev, 2006. 19(3): p. 531–45.

13. Andersen, K.G., et al., The proximal origin of SARS-CoV-2. Nat Med, 2020. 26(4): p. 450–452.

14. Brook, C.E. and A.P. Dobson, Bats as ‘special’ reservoirs for emerging zoonotic pathogens. Trends Microbiol, 2015. 23(3): p. 172–80.

15. Mollentze, N. and D.G. Streicker, Viral zoonotic risk is homogenous among taxonomic orders of mammalian and avian reservoir hosts. Proceedings of the National Academy of Sciences, 2020. 117(17): p. 9423–9430.

16. Irving, A.T., et al., Lessons from the host defences of bats, a unique viral reservoir. Nature, 2021. 589(7842): p. 363–370.

17. Pavlovich, S.S., et al., The Egyptian Rousette Genome Reveals Unexpected Features of Bat Antiviral Immunity. Cell, 2018. 173(5): p. 1098-1110.e18.

18. Moreno Santillán, D.D., et al., Large-scale genome sampling reveals unique immunity and metabolic adaptations in bats. Mol Ecol, 2021. 30(23): p. 6449–6467.

19. O’Connor, E.A., et al., Wetter climates select for higher immune gene diversity in resident, but not migratory, songbirds. Proceedings of the Royal Society B: Biological Sciences, 2020. 287(1919): p. 20192675.

20. Westerdahl, H., et al., The genomic architecture of the passerine MHC region: High repeat content and contrasting evolutionary histories of single copy and tandemly duplicated MHC genes. Molecular Ecology Resources, 2022. 22(6): p. 2379–2395.

21. Murphy, K., et al., Janeway’s Immunobiology. 9th ed. 2016, New York: Garland Science.

22. Nowak, M.A., K. Tarczy-Hornoch, and J.M. Austyn, The optimal number of major histocompatibility complex molecules in an individual. Proc Natl Acad Sci U S A, 1992. 89(22): p. 10896–9.

23. Woelfing, B., et al., Does intra-individual major histocompatibility complex diversity keep a golden mean? Philosophical transactions of the Royal Society of London. Series B, Biological sciences, 2009. 364(1513): p. 117–128.

24. Lenz, T.L., Computational prediction of MHC II-antigen binding supports divergent allele advantage and explains trans-species polymorphism. Evolution, 2011. 65(8): p. 2380–90.

25. Ng, J.H., et al., Evolution and comparative analysis of the bat MHC-I region. Sci Rep, 2016. 6: p. 21256.

26. Lu, D., et al., Peptide presentation by bat MHC class I provides new insight into the antiviral immunity of bats. PLoS Biol, 2019. 17(9): p. e3000436.

27. Qu, Z., et al., Structure and Peptidome of the Bat MHC Class I Molecule Reveal a Novel Mechanism Leading to High-Affinity Peptide Binding. J Immunol, 2019. 202(12): p. 3493–3506.

28. O’Connor, E. and H. Westerdahl, Trade-offs in expressed major histocompatibility complex diversity seen on a macroevolutionary scale among songbirds. Evolution, 2021. 75(5): p. 1061–1069.

29. Chappell, P., et al. Expression levels of MHC class I molecules are inversely correlated with promiscuity of peptide binding. eLife, 2015. 4, e05345 DOI: 10.7554/elife.05345.

30. Koleček, J., et al., Cross-continental migratory connectivity and spatiotemporal migratory patterns in the great reed warbler. Journal of Avian Biology, 2016. 47(6): p. 756–767.

31. Westerdahl, H., H. Wittzell, and T. von Schantz, Polymorphism and transcription of Mhc class I genes in a passerine bird, the great reed warbler. Immunogenetics, 1999. 49(3): p. 158–170.

32. Westerdahl, H., H. Wittzell, and T. von Schantz, Mhc diversity in two passerine birds: no evidence for a minimal essential Mhc. Immunogenetics, 2000. 52(1): p. 92–100.

33. Westerdahl, H., et al., MHC class I typing in a songbird with numerous loci and high polymorphism using motif-specific PCR and DGGE. Heredity (Edinb), 2004. 92(6): p. 534–42.

34. Rasmussen, M., et al., Uncovering the peptide-binding specificities of HLA-C: a general strategy to determine the specificity of any MHC class I molecule. J Immunol, 2014. 193(10): p. 4790–802.

35. Saper, M.A., P.J. Bjorkman, and D.C. Wiley, Refined structure of the human histocompatibility antigen HLA-A2 at 2.6 Å resolution. Journal of Molecular Biology, 1991. 219(2): p. 277–319.

36. Rasmussen, M., et al., Large scale analysis of peptide-HLA-I stability, 2014.

37. Harndahl, M., et al., Large scale analysis of peptide-HLA class I interactions, 2007.

38. Harndahl, M., et al., Large scale analysis of peptide-HLA class I interactions, 2006.

39. Harndahl, M., et al., Large scale analysis of peptide-HLA class I interactions, 2010.

40. Thomsen, M., et al., MHCcluster, a method for functional clustering of MHC molecules. Immunogenetics, 2013. 65(9): p. 655–665.

41. Sidney, J., et al., HLA class I supertypes: a revised and updated classification. BMC immunology, 2008. 9: p. 1–1.

42. Sjöberg, S., et al., Extreme altitudes during diurnal flights in a nocturnal songbird migrant. Science, 2021. 372(6542): p. 646–648.

43. O’Connor, E., et al., The evolution of immunity in relation to colonization and migration. Nature Ecology & Evolution, 2018. 2: p. 841–849.

44. Wei, X., et al., Structure and Peptidomes of Swine MHC Class I with Long Peptides Reveal the Cross-Species Characteristics of the Novel N-Terminal Extension Presentation Mode. J Immunol, 2022. 208(2): p. 480–491.

45. Barik, S., The Uniqueness of Tryptophan in Biology: Properties, Metabolism, Interactions and Localization in Proteins. International Journal of Molecular Sciences, 2020. 21(22): p. 8776.

46. Kaleta, C., et al., Metabolic costs of amino acid and protein production in Escherichia coli.

47. Cedano, J., et al., Relation between amino acid composition and cellular location of proteins. J Mol Biol, 1997. 266(3): p. 594–600.

48. Du, M.Z., et al., The GC Content as a Main Factor Shaping the Amino Acid Usage During Bacterial Evolution Process. Front Microbiol, 2018. 9: p. 2948.

49. Hoyle, L. and S.P. Davies, Amino acid composition of the protein components of influenza virus A. Virology, 1961. 13(1): p. 53–57.

50. Moura, A., M.A. Savageau, and R. Alves, Relative amino acid composition signatures of organisms and environments. PLoS One, 2013. 8(10): p. e77319.

51. Akashi, H. and T. Gojobori, Metabolic efficiency and amino acid composition in the proteomes of Escherichia coli and Bacillus subtilis. 2002.

52. Creixell, P., et al., Mutational properties of amino acid residues: implications for evolvability of phosphorylatable residues. Philosophical transactions of the Royal Society of London. Series B, Biological sciences, 2012. 367(1602): p. 2584–2593.

53. Yampolsky, L.Y. and A. Stoltzfus, The exchangeability of amino acids in proteins. Genetics, 2005. 170(4): p. 1459–1472.

54. Norn, C., I. André, and D.L. Theobald, A thermodynamic model of protein structure evolution explains empirical amino acid substitution matrices. Protein Sci, 2021. 30(10): p. 2057–2068.

55. Zhang, X.H., D. Baronas-Lowell, and H. Weissbach, Unique metabolic roles of methionine both free and in proteins. Current Topics in Peptide and Protein Research, 2011. 12: p. 1–15.

56. Brosnan, J.T., et al., Methionine: A metabolically unique amino acid. Livestock Science, 2007. 112(1): p. 2–7.

57. Aledo, J.C., Methionine in proteins: The Cinderella of the proteinogenic amino acids. Protein Science, 2019. 28(10): p. 1785–1796.

58. Lim, J.M., G. Kim, and R.L. Levine, Methionine in Proteins: It’s Not Just for Protein Initiation Anymore. Neurochem Res, 2019. 44(1): p. 247–257.

59. Bohórquez, H.J., C.F. Suárez, and M.E. Patarroyo, Mass & secondary structure propensity of amino acids explain their mutability and evolutionary replacements. Scientific Reports, 2017. 7(1): p. 7717.

60. Pymm, P., et al., MHC-I peptides get out of the groove and enable a novel mechanism of HIV-1 escape. Nat Struct Mol Biol, 2017. 24(4): p. 387–394.

61. Li, X., et al., Crystal structure of HLA-B*5801 with a TW10 HIV Gag epitope reveals a novel mode of peptide presentation. Cell Mol Immunol. 2017 Jul;14(7):631–634. doi: 10.1038/cmi.2017.24. Epub 2017 May 29.

62. Ohta, Y. and M.F. Flajnik, Coevolution of MHC genes (LMP/TAP/class Ia, NKT-class Ib, NKp30-B7H6): lessons from cold-blooded vertebrates. Immunol Rev, 2015. 267(1): p. 6–15.

63. Yoder, J.A. and G.W. Litman, The phylogenetic origins of natural killer receptors and recognition: relationships, possibilities, and realities. Immunogenetics, 2011. 63(3): p. 123–41.

64. Straub, C., et al., Chicken NK cell receptors. Dev Comp Immunol, 2013. 41(3): p. 324–33.

65. Kaufman, J., From Chickens to Humans: The Importance of Peptide Repertoires for MHC Class I Alleles. Front Immunol, 2020. 11: p. 601089.

66. Brackenridge, S., et al., An Early HIV Mutation within an HLA-B*57-Restricted T Cell Epitope Abrogates Binding to the Killer Inhibitory Receptor 3DL1. Journal of Virology, 2011. 85(11): p. 5415–5422.

67. Follin, E., et al., In silico peptide-binding predictions of passerine MHC class I reveal similarities across distantly related species, suggesting convergence on the level of protein function. Immunogenetics, 2013. 65(4): p. 299–311.

68. Lundberg, M., et al., Characterisation of a transcriptome to find sequence differences between two differentially migrating subspecies of the willow warbler Phylloscopus trochilus. Bmc Genomics, 2013. 14.

69. Ekblom, R., et al., Characterization of the house sparrow (Passer domesticus) transcriptome: a resource for molecular ecology and immunogenetics. Molecular ecology resources, 2014. 14(3): p. 636–46.

70. Wada, K., et al., A molecular neuroethological approach for identifying and characterizing a cascade of behaviorally regulated genes. Proceedings of the National Academy of Sciences of the United States of America, 2006. 103(41): p. 15212–7.

71. Strandh, M., et al., Characterization of MHC class I and II genes in a subantarctic seabird, the blue petrel, Halobaena caerulea (Procellariiformes). Immunogenetics, 2011. 63(10): p. 653–666.

72. Østergaard Pedersen, L., et al., Efficient assembly of recombinant major histocompatibility complex class I molecules with preformed disulfide bonds. European Journal of Immunology, 2001. 31(10): p. 2986–2996.

73. Stryhn, A., et al., Peptide binding specificity of major histocompatibility complex class I resolved into an array of apparently independent subspecificities: quantitation by peptide libraries and improved prediction of binding. Eur J Immunol, 1996. 26(8): p. 1911–8.

74. Hansen, A.M., et al., Characterization of binding specificities of bovine leucocyte class I molecules: impacts for rational epitope discovery. Immunogenetics, 2014. 66(12): p. 705–718.

75. Thomsen, M.C.F. and M. Nielsen, Seq2Logo: a method for construction and visualization of amino acid binding motifs and sequence profiles including sequence weighting, pseudo counts and two-sided representation of amino acid enrichment and depletion. Nucleic acids research, 2012. 40(Web Server issue): p. W281–W287.

76. Nielsen, M., O. Lund, and C. Lundegaard. MHCcluster, a method for functional clustering of MHC molecules. in ISCB-Latin 2012.

77. Jurtz, V., et al., NetMHCpan-4.0: Improved Peptide-MHC Class I Interaction Predictions Integrating Eluted Ligand and Peptide Binding Affinity Data. J Immunol, 2017. 199(9): p. 3360–3368.

78. Nagai, K. and H.C. Thogersen, Synthesis and sequence-specific proteolysis of hybrid proteins produced in Escherichia coli. Methods Enzymol, 1987. 153: p. 461–81.

79. Garboczi, D.N., D.T. Hung, and D.C. Wiley, HLA-A2-peptide complexes: refolding and crystallization of molecules expressed in Escherichia coli and complexed with single antigenic peptides. Proceedings of the National Academy of Sciences of the United States of America, 1992. 89(8): p. 3429–3433.

80. Kabsch, W., XDS. Acta Crystallogr D Biol Crystallogr, 2010. 66(Pt 2): p. 125–32.

81. Winn, M.D., et al., Overview of the CCP4 suite and current developments. Acta Crystallogr D Biol Crystallogr, 2011. 67(Pt 4): p. 235–42.

82. McCoy, A.J., et al., Phaser crystallographic software. Journal of Applied Crystallography, 2007. 40(4): p. 658–674.

83. Murshudov, G.N., A.A. Vagin, and E.J. Dodson, Refinement of Macromolecular Structures by the Maximum-Likelihood Method. Acta Crystallographica Section D, 1997. 53(3): p. 240–255.

84. Liebschner, D., et al., Macromolecular structure determination using X-rays, neutrons and electrons: recent developments in Phenix. Acta Crystallographica Section D, 2019. 75(10): p. 861–877.

85. Emsley, P. and K. Cowtan, Coot: model-building tools for molecular graphics. Acta Crystallogr D Biol Crystallogr, 2004. 60(Pt 12 Pt 1): p. 2126–32.

86. Krissinel, E. and K. Henrick, Secondary-structure matching (SSM), a new tool for fast protein structure alignment in three dimensions. Acta Crystallogr D Biol Crystallogr, 2004. 60(Pt 12 Pt 1): p. 2256–68.

87. Thomsen, M. C., & Nielsen, M. (2012). Seq2Logo: a method for construction and visualization of amino acid binding motifs and sequence profiles including sequence weighting, pseudo counts and two-sided representation of amino acid enrichment and depletion. Nucleic acids research. doi:10.1093/nar/gks469

