## Supplementary Material for "Extraordinary peptide-binding mode of a songbird MHC class-I molecule suggests mechanism to counter pathogen immune evasion"

**Figure S1**

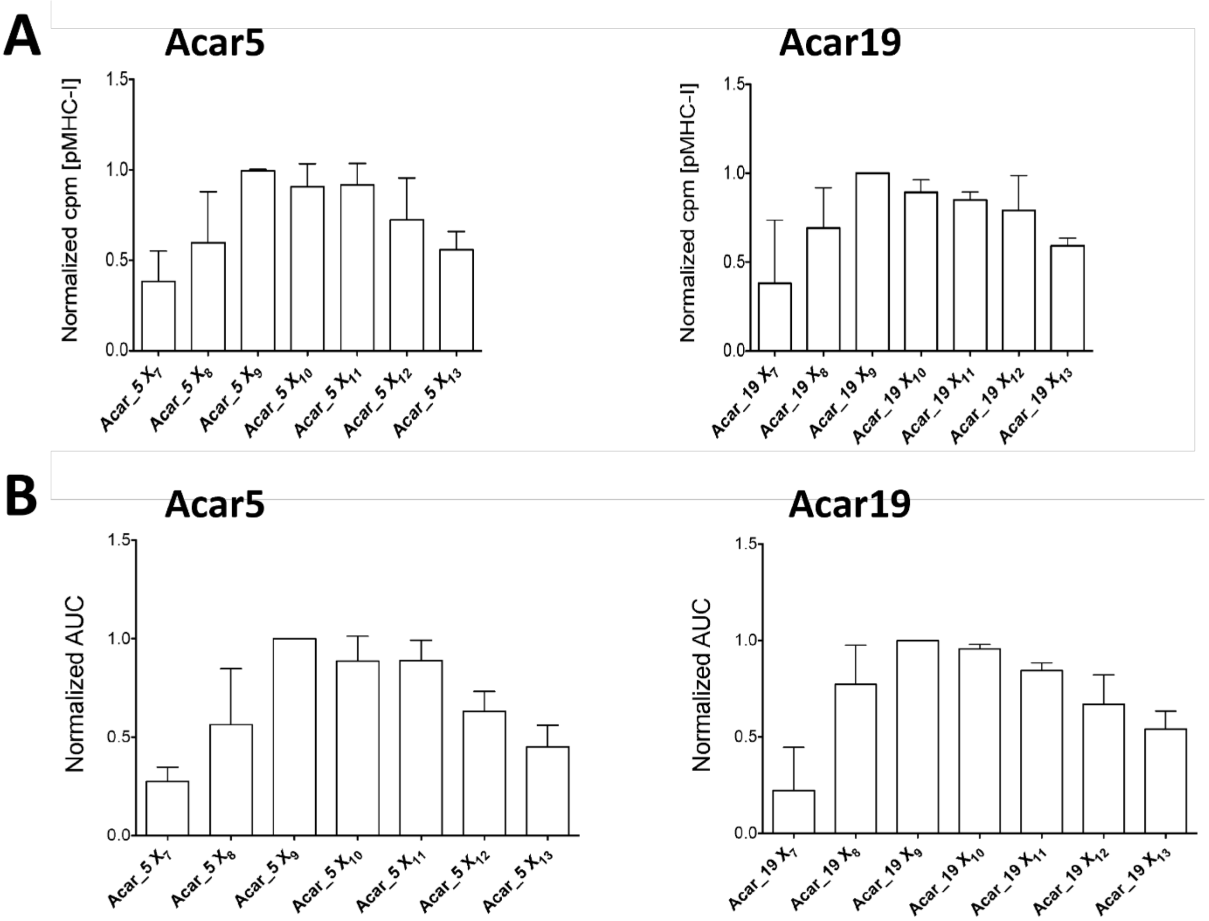

**Figure S1: Great reed warbler MHC-I molecules show classical MHC-I peptide-length preferences.** The heavy chains (hc) of Acar5 (left) and Acar19 (right) that are closely related to Acar3 (95.3 % and 88.7 % sequence identity, respectively) were folded in the presence of $X_7$-$X_{13}$ peptide libraries with $^{125}$I-labeled *Acar* $\beta_2$m. **(A)** The number of complexes formed after 24 hours folding at 18 °C was analysed in a scintillation proximity-based assay (SPA) at 37 °C. Acar5 and Acar19 formed significantly less complexes with 7-mer peptides compared to 9-, 10- and 11-mers. **(B)** The dissociation of Acar pMHC-I complexes formed with $X_7$-$X_{13}$ peptide libraries was monitored for 24 hours to investigate complex stability. The normalised area under the curve (AUC) was calculated for each Acar-$X_n$ MHC-I allomorph combination and values were compared to Acar-$X_9$ MHC-I, which formed the most stable complexes with both, Acar5 and Acar19.

**Figure S2**

| | | Peptide position | | | | | | | | |
|---|---|---|---|---|---|---|---|---|---|---|
| | | 1 | 2 | 3 | 4 | 5 | 6 | 7 | 8 | 9 |
| Amino acid | **A** | 0,9 | 0,8 | 0,9 | 1,0 | 1,4 | 1,3 | 1,1 | 1,0 | 0,8 |
| | **C** | 0,9 | 1,0 | 0,7 | 0,7 | 1,2 | 1,0 | 1,0 | 1,0 | 1,0 |
| | **D** | 1,2 | 1,0 | 1,1 | 0,7 | 0,7 | 1,1 | 0,9 | 1,4 | 1,3 |
| | **E** | 1,1 | 0,7 | 0,8 | 1,1 | 0,9 | 1,3 | 1,3 | 1,3 | 1,2 |
| | **F** | 0,9 | 1,1 | *0,3* | 1,0 | 1,0 | 1,0 | 1,1 | 1,0 | **2,6** |
| | **G** | 1,1 | 0,6 | 0,6 | 1,0 | 1,0 | 0,7 | 0,8 | 0,7 | 1,1 |
| | **H** | 0,7 | 0,8 | 0,8 | 0,8 | 0,9 | 1,1 | 1,2 | 1,2 | 0,8 |
| | **I** | 0,9 | 1,0 | 1,0 | 0,9 | 1,1 | *0,5* | 0,8 | 0,8 | 1,5 |
| | **K** | 1,1 | 1,4 | 1,2 | 1,0 | 1,3 | 1,0 | 1,1 | 0,8 | *0,3* |
| | **L** | 1,0 | 0,7 | 1,2 | 1,1 | 1,1 | 1,0 | 1,0 | 1,1 | 1,9 |
| | **M** | 0,9 | 1,6 | **2,1** | 1,3 | 1,3 | 1,0 | 0,9 | 1,0 | 1,5 |
| | **N** | 1,2 | 0,7 | 1,2 | 1,2 | 0,8 | 1,0 | 1,2 | 1,0 | 0,6 |
| | **P** | 0,8 | 1,0 | 1,0 | 1,2 | 1,3 | 1,8 | 1,4 | 1,3 | 0,6 |
| | **Q** | 1,2 | 1,0 | 1,0 | 1,2 | 1,1 | 1,2 | 1,2 | 0,9 | *0,4* |
| | **R** | 0,9 | 1,3 | 0,9 | 1,1 | 0,7 | 1,0 | 1,1 | 0,7 | 0,6 |
| | **S** | 1,0 | 0,6 | 1,0 | 1,0 | 0,7 | 0,6 | 1,0 | 0,9 | *0,4* |
| | **T** | 0,9 | 1,2 | 1,2 | 1,0 | 1,1 | 1,1 | 0,9 | 1,0 | *0,4* |
| | **V** | 1,1 | 1,1 | 1,1 | 0,8 | 1,0 | 1,2 | 1,0 | 1,0 | 1,2 |
| | **W** | 1,1 | 1,2 | 1,4 | 1,3 | 0,7 | 1,0 | 1,0 | 0,8 | 0,9 |
| | **Y** | 1,2 | 0,9 | 1,1 | 0,8 | 0,8 | 1,0 | 1,1 | 1,0 | 0,7 |
| | **Sum** | **20** | **20** | **20** | **20** | **20** | **20** | **20** | **20** | **20** |
| | **AP** | 0,6 | 1,5 | 2,5 | 0,6 | 1,2 | 1,5 | 0,5 | 0,7 | 6,7 |

**Figure S2: PSCPL matrix for nonameric peptides binding to Acar3.** The matrix reflects the peptide-binding properties of Acar3 through the relative binding (RB) value for each amino acid at each position of nonameric peptides from the PSCPL. An RB-value of ≥ 2 defines a favoured amino acid at a specific position, while an RB-value of ≤ 0.5 defines a disfavoured amino acid at a specific position. The anchor-position (AP) value defines the relative contribution of each position to peptide binding (Rasmussen *et al.*, 2014). The logo of the PSCPL-derived Acar3 binding motif ***(Figure 1)*** was made using the Seq2Logo 1.1 (Thomsen & Nielsen, 2012) server as P-Weighted Kullback-Leibler logo.

**Figure S3**

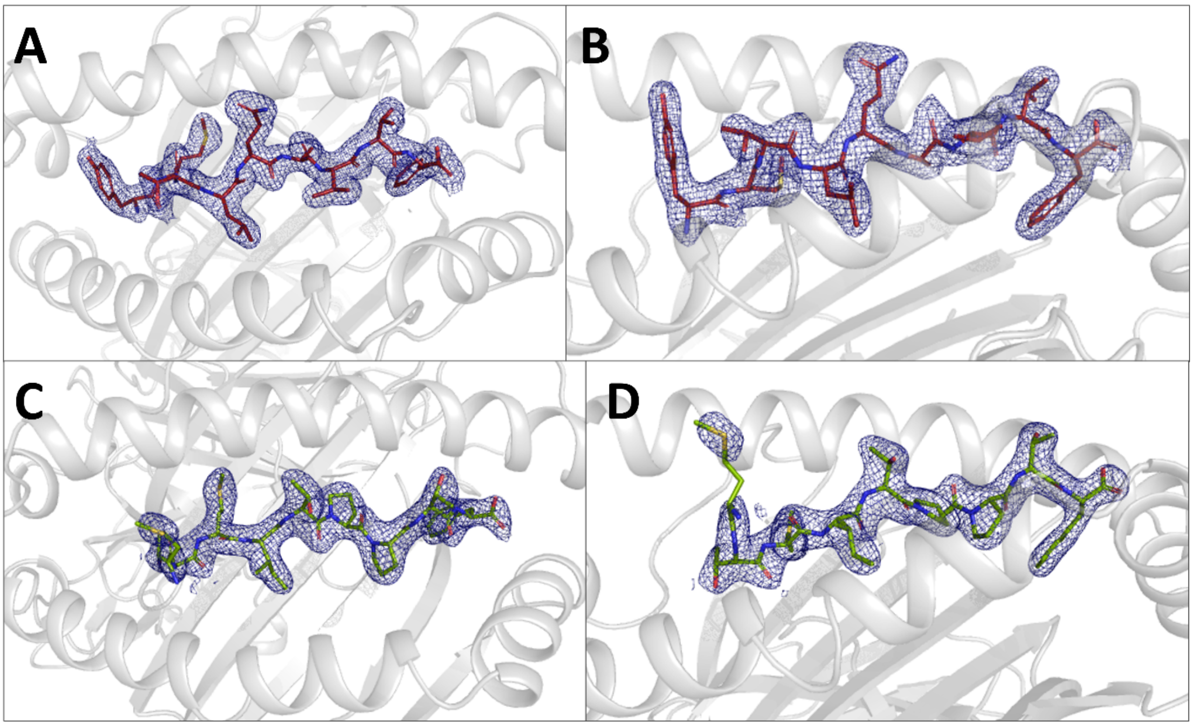

**Figure S3: Electron density maps for P2 and P3.** Top **(A)** and side **(B)** view of the 2Fo-Fc electron density map of P2 (red, stick representation) shown as blue mesh at a contour level of 1.0 σ. Top **(C)** and side **(D)** view of the 2Fo-Fc electron density map of P3 (green, stick representation) shown as blue mesh at a contour level of 1.0 σ. The Acar3 PBG is shown as cartoon in grey.

**Figure S4**

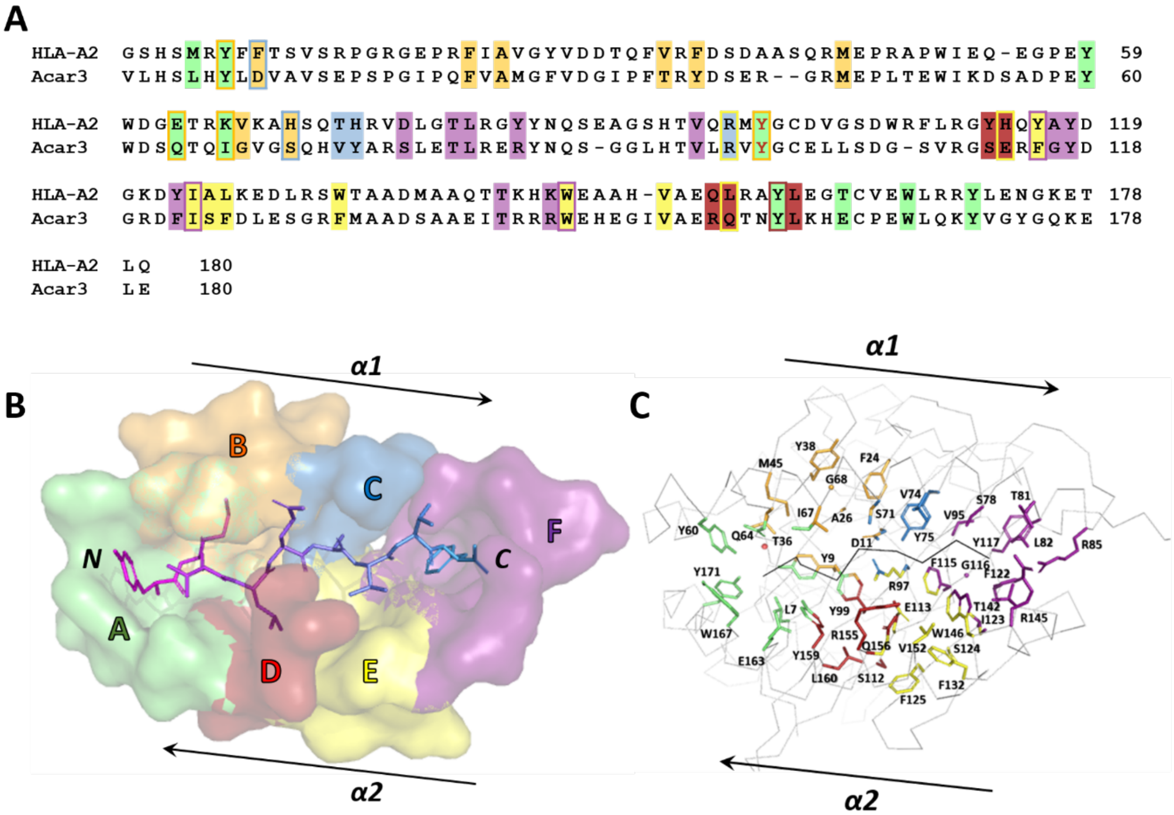

**Figure S4: The six pockets of the Peptide Binding Groove (PBG) common among HLA-A2 and Acar3. (A)** Alignment of the α1 and α2 domains of the HLA-A2 molecule as described in (Saper *et al.*, 1991) (PDB code 3HLA, aa 1-180) and Acar3 (aa 3-180). The residues that participate in the formation of the six pockets of the PBG are highlighted in different colours. The residues are numbered according to the Acar3 sequence and the HLA-A2 sequence (in parentheses): **Pocket A** (green): L7 (M5), Y9 (Y7), Y60 (Y59), Q64 (E63), I67 (K66), Y99 (Y99), Y159 (Y159), E163 (T163), W167 (W167), and Y171 (Y171); **Pocket B** (orange): Y9 (Y7), D11 (F9), A26 (A24), T36 (V34), M45 (M45), Q64 (E63), I67 (K66), G68 (V67), S71 (H70) and Y99 (Y99). In Acar3, F24 (F22) and Y38 (F36) additionally participate in creating this pocket. **Pocket C** (blue): D11 (F9), S71(H70), V74 (T73), Y75 (H74) and R97 (R97); **Pocket D** (red): Y99 (Y99), the backbone of S112 (Y113); E113 (H114), R155 (Q155), Q156 (L156), Y159 (Y159) and L160 (L160); **Pocket E** (yellow): R97 (R97), E113 (H114), F132 (W133), W146 (W147), V152 (V152) and Q156 (L156). In Acar3, F115 (Y116), I123 (I124) and the backbone atoms of S124 (A125) and F125 (L126) additionally contribute to the pocket surface. **Pocket F** (magenta): S78 (D77), T81 (T80), L82 (L81), R85 (Y84), F115 (Y116), F122 (Y123), T142 (T143), R145 (K146) and W146 (W147). In Acar3, V95 (V95), the backbone atoms of G116 (A117) and Y117 (Y118), and I123 (I124) additionally participate in creating this pocket. Residues that are part of two pockets contain a background and a frame in two different colours according to the colouring scheme of the corresponding pockets. In case a residue belongs to three pockets, the aa letter is coloured corresponding to the third pocket. **(B)** Top view onto the PBG of the Acar3-P2 complex with the pockets as surface representation coloured according to **(A)** and P2 shown in stick representation. **(C)** Stick representation of the pockets' residues coloured according to **(A)**. The Acar3 scaffold and P2 are shown as grey and black ribbon, respectively. **Pocket A** is open towards the solvent and creates an anchor for the N-termini of the bound peptides. A conserved water molecule (**(C)**, red sphere) is coordinated by Y9, T36, Y60 and Q64 and supports the specific alignment of those residues. The surface potential of the pocket can be altered by the Q64-side chain adapting different orientations. **Pocket B** adopts an elongated shape and has an increased depth and volume compared to pocket B in HLA-A2, which is due to the smaller residues at positions 11, 68 and 71. The pocket hence permits accommodation of bulky residues and harbours the methionine-anchor residue of P2 and P3 (M2 in P2, M3 in P3). Since the M-side chain does not completely fill the cavity, the residual space is occupied by one water molecule in the P3 structure, and two water molecules in the Acar3-P2 complex. The bottom of **pocket C** is formed by the aromatic portion of Y75. S71 and the guanidinium group of R97 create a rim that segregates pocket C from pocket B. According to the shallow and neutral character of pocket C, short and hydrophobic residues, such as A (P2) or P (P3) at position 6 are advantageous for peptide binding. **Pocket D** is separated from the adjacent pocket A by the shared side chain of Y159. This pocket shows different morphologies and surface potentials in the two structures and can likely accommodate amino acids of different shapes and polarities. Residues E113, R155 and Q156 have crucial impact on the shape and surface potential of pocket D and their side-chain conformations vary among both structures. In the Acar3-P2 complex, R155 can be present two different conformations either directed towards P2 or towards the solvent. **Pocket E** shows some extent of variability between the two structures with surface potential ranging from neutral (P2) to slightly negative (P3). The cavity has a round shape and varies in depth and width depending on the orientation of residues E113, F115, F132, W146 and Q156. Moreover, water molecules can be present in the cavity to mediate hydrogen bonds between the peptide and the PBG residues. **Pocket F** shows a highly positive surface potential around its upper rim as well as clearly positively polarised surface characteristics within the lower parts of the cavity, which is of importance to anchor the C-terminus of the antigenic peptide. The strong polarisation around the rim is primarily achieved through contributions from the guanidinium groups of R85 and R145, as well as W146, whereas the central part and the bottom of the cavity comprise exclusively hydrophobic/aromatic residues.

**Figure S5**

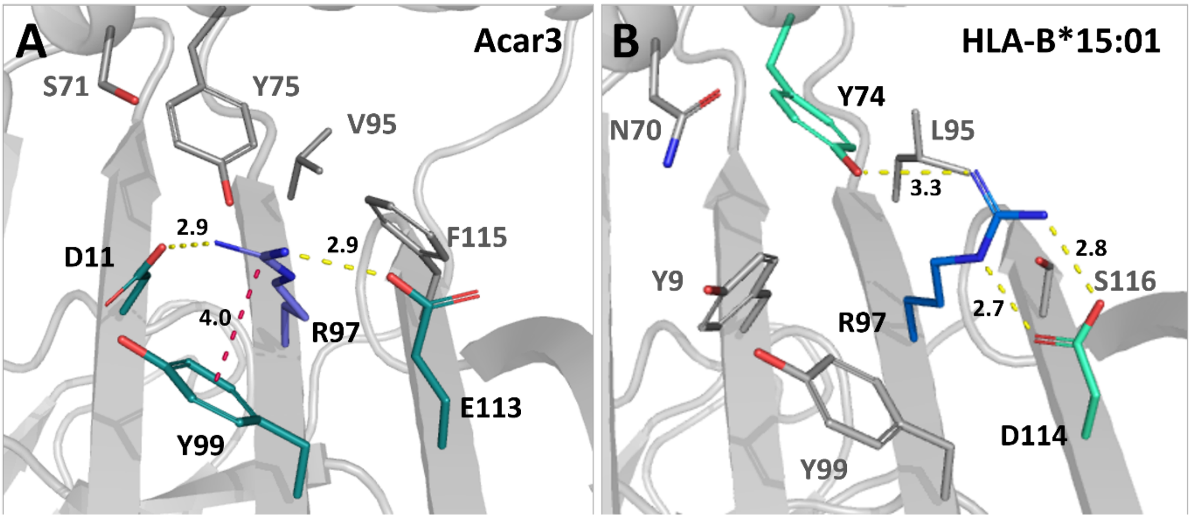

**Figure S5: Major interactions of Arg97 in Acar3 and HLA-B*15:01. (A)** In Acar3, Arg97 is oriented towards the centre of the PBG, thereby establishing close interactions with Asp11 lining pockets B and C, with Glu113 which is part of pockets D and E, and with Tyr99 flanking pockets A, B and D. **(B)** In HLA-B*15:01, Arg97 is oriented towards the area of the PBG that accommodates the C-terminal residues of bound peptides, forming hydrogen bonds with Tyr74 and Asp114 which belong to pockets C and D/E, respectively. The closest neighbours of Arg97 (blue sticks) in the PBG are shown as sticks; hydrogen-bonding and cation-π interaction partners are highlighted in teal **(A)** and greencyan **(B)**. Hydrogen bonds are indicated as yellow, dashed lines; cation-π interactions are depicted as red, dashed lines. Distances are given in Å.

**Figure S6**

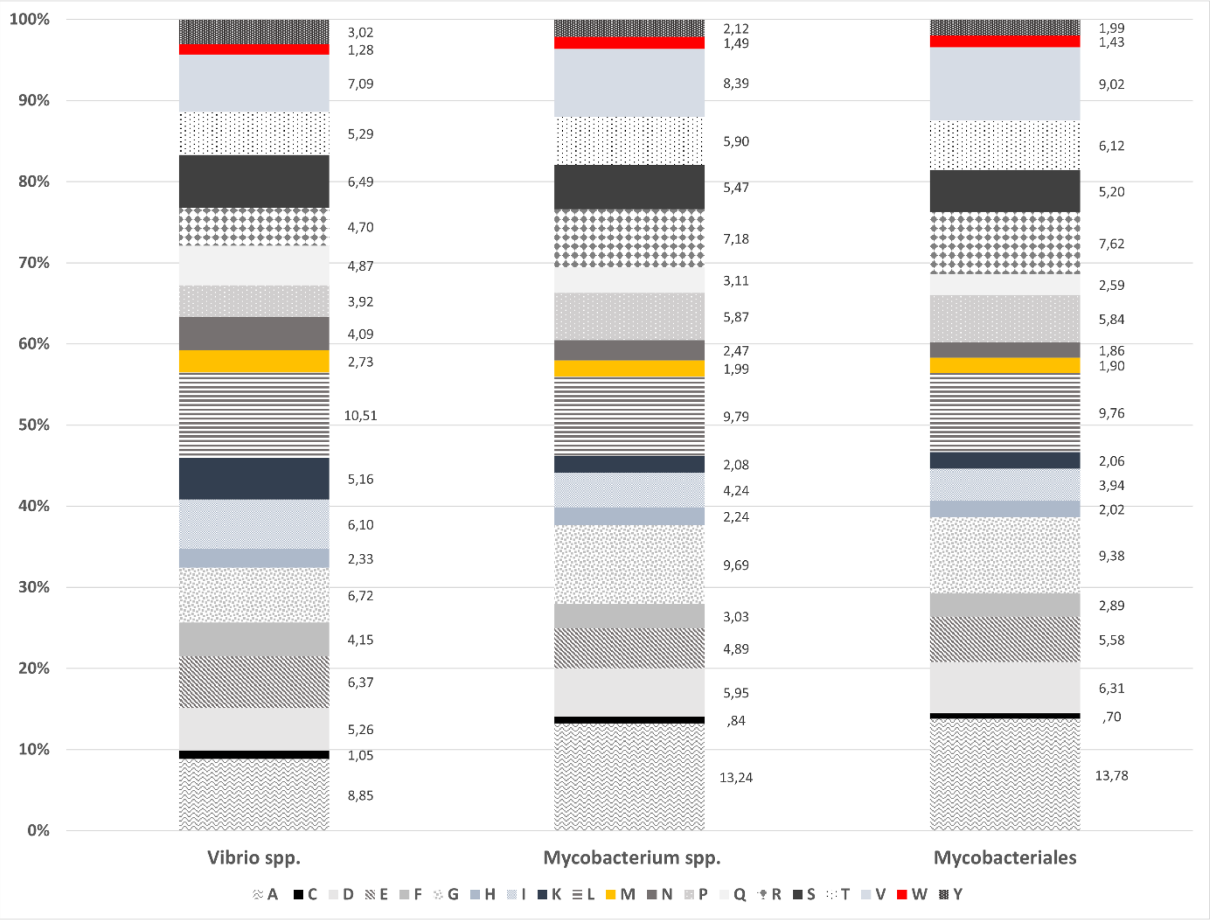

**Figure S6: Percentages of amino acids within the proteomes of bacteria representing potential antigen sources for Acar3 binding.** Proteomes were retrieved from UniProt (Consortium, 2020). *Vibrio spp.* comprises the proteomes of *V. cholerae* (UP000000584), *V. vulnificus* (UP000002675) and *V. parahaemolyticus* (UP000002493); *Mycobacterium spp.* includes the proteomes of *M. tuberculosis* (UP000001584), *M. paraense* (UP000193285), *M. marinum* (UP000001190), *M. canettii* (UP000008896), *M. simiae* (UP000193040), *M. gordonae* (UP000193928) and *M. malmoense* (UP000243140); and Mycobacteriales contains the proteomes of *Dietzia* and *Tsukamurella spp., i.e. D. cinnamea* (UP000295805), *T. sputi* (UP000319792) and *T. paurometabola* (UP000271626). Amino acids (A-Y) are given in one-letter code.

**Table S1: Stability of Acar3 in complex with the peptides obtained through PSCPL analysis**

| Sequence | Stability T $^1/_2$ (h) | Sequence | Stability T $^1/_2$ (h) |
|---|---|---|---|
| AMSAQAAAF (P1) | 11.3 | RMMETWHPL | 0.9 |
| YMTLQAVTF (P2) | 11.2 | MMQVWIQPL | 0.9 |
| MTMITPPTF (P3) | 10.3 | FHMSKPSDL | 0.8 |
| KTMMQAHDL | 9.9 | LMMNGTSAM | 0.8 |
| QWALLPYLF | 9.9 | MMWATAQAL | 0.8 |
| GMVGAEFSL | 7.6 | YWMGGTTYF | 0.8 |
| RYMSKTYNF | 7.5 | IYLPIVHPF | 0.7 |
| MMMSTAVAF | 7.3 | GPMKLVMAF | 0.7 |
| MMMPMFNAF | 7.2 | WQLTSIWPI | 0.7 |
| RRMATTFTF | 6.4 | YPMSIPATL | 0.7 |
| IFLKPEETF | 6.1 | FQMGGIGPM | 0.7 |
| KAMSTPFSL | 5.3 | REMINHYQV | 0.7 |
| AEMRETHWL | 5.0 | AVMFFPFWF | 0.6 |
| LTMNLVSDI | 4.5 | QMSWLPLCV | 0.6 |
| MMHASTSPF | 4.2 | LDMNVVKEL | 0.6 |
| KQMSQPYAV | 4.0 | RQNAAIEAL | 0.6 |
| AMPKTIYEL | 3.8 | GAMYALANF | 0.6 |
| YQVKYVSPV | 3.6 | GRLVTVNPF | 0.5 |
| MPMSMPIPM | 3.2 | RMAMIPRTL | 0.5 |
| WMMAMRYPI | 3.1 | VMMAKRPLI | 0.5 |
| SPMETTAEF | 3.0 | YLQQNTHTL | 0.5 |
| ETMKPAAMV | 2.7 | LRWLAKSFF | 0.5 |
| YMWECPDFF | 2.6 | QQMFPGAPF | 0.4 |
| LQMLGENVL | 2.6 | KAMRPWQSF | 0.4 |
| KIMQVDRPM | 2.6 | KMAAFPETL | 0.4 |
| AKIEAPLLL | 2.5 | IRMEFQSVF | 0.4 |
| YLYLRPYAL | 2.4 | KLMDVVYSI | 0.4 |
| SLAIDAYPL | 2.1 | YMWLGARYL | 0.4 |
| KMNAKAATL | 2.1 | LMDENTYAM | 0.4 |
| VMPLLVWLF | 1.9 | QQLEADYTF | 0.4 |
| REWSGLNYL | 1.9 | YLLEMLWRL | 0.4 |
| MTYKAAFDL | 1.8 | KTAAAAAAF | 0.4 |
| RRWIAPHPL | 1.7 | RMMATKDSF | 0.4 |
| RMLPKLAEF | 1.6 | LPLPAQQPF | 0.3 |
| SVYFAAFAF | 1.6 | GMNVTAPAL | 0.3 |
| RATQPAAEF | 1.6 | SQMGLSCAL | 0.3 |
| QAYAAPQLF | 1.5 | VIMWYNYLF | 0.3 |
| GLAATSFPL | 1.5 | LPRWPPPQL | 0.3 |
| VPMEKLKAL | 1.4 | FRMLAWHVL | 0.3 |
| IMDASSFTL | 1.4 | FMIVNNFPV | 0.3 |
| KQLESVMYL | 1.3 | AMAAGLSSL | 0.3 |
| FQANMFLTL | 1.2 | AMMWRIAQL | 0.3 |
| FLLENAAYL | 1.2 | FLDKGTYTL | 0.3 |
| HELSLFWPL | 1.1 | STEFIPNLF | 0.3 |
| YMHGSIHEV | 0.9 | ILMEHIHKL | 0.2 |
| SMMPEAMTI | 0.9 | DYQGLPLEF | 0.2 |
| WMACNSAAF | 0.9 | SYPPPPASF | 0.2 |

**Table S2: Interactions of P2 within the Acar3 PBG**

| P2 | | | Acar3 | | |
|---|---|---|---|---|---|
| ***Hydrogen bonds (≤ 3.4 Å)*** | | | | | |
| *Residue* | *Atom* | *Distance (Å)* | *Residue* | *Atom* | *Pocket* |
| $Y_1$ | N | 2.7 | Y171 | Oη | A |
| | N | 2.7 | Y9 | Oη | |
| | Oη | 2.7 | S63 | Oγ | |
| | O | 2.6 | Y159 | Oη | |
| $M_2$ | N | 3.3 | Q64 | Oε1 | B |
| | N | 3.2 | Y9 | Oη | |
| $T_3$ | Oγ1 | 3.2 | E163 | Oε2 | - |
| $L_4$ | O | 3.0 | R97 | Nη1 | D |
| | O | 2.8 | R97 | Nη2 | |
| $Q_5$ | N | 3.4 | R155 | Nη2, A | - |
| | O | 2.3 | R155 | Nη2, A | |
| $A_6$ | N | 2.8 | S71 | Oγ | C |
| $V_7$ | - | - | - | - | E |
| $T_8$ | Oγ1 | 3.0 | S78 | Oγ | - |
| | O | 3.1 | W146 | Nε1 | |
| $F_9$ | N | 3.0 | S78 | Oγ | F |
| | O | 2.6 | T142 | Oγ1 | |
| | O | 3.2 | R145 | Nε | |
| | O | 2.8 | R85 | Nη2 | |
| | OXT | 3.3 | R145 | Nη2 | |
| ***Water-bridged hydrogen bonds (≤ 3.4 Å)*** | | | | | |
| *Residue (P2)* | *Atom* | *Distance (Å)* P2-residue – $H_2O$ | *Distance (Å)* $H_2O$ – Acar3-residue | *Residue (Acar3)* | *Atom* |
| $V_7$ | N | 3.2 | 2.8 | E113 | Oε2 |
| | | | 2.8 | Q156 | Nε2 |
| | O | 2.8 | 2.7 | S78 | Oγ |
| $T_8$ | Oγ1 | 2.7 | 2.8 | T81 | Oγ1 |
| $F_9$ | OXT | 2.7 | 2.8 | T81 | Oγ1 |
| ***Salt bridges (≤ 4.0 Å)*** | | | | | |
| *Residue* | *Atom* | *Distance (Å)* | *Residue* | *Atom* | *Pocket* |
| $F_9$ | O | 3.9 | R85 | Nη1 | F |

| | | | | | |
|---|---|---|---|---|---|
| | | 2.8 | | Nη2 | |
| | | 3.2 | R145 | Nε | |
| | | 3.9 | | Nη2 | |
| | OXT | 4.0 | R145 | Nε | |
| | | 3.3 | | Nη2 | |
| ***Hydrophobic contacts/π-stacking (≤ 4.0 Å)*** | | | | | |
| *Residue* | *Atom* | *Distance (Å)* | *Residue* | *Atom* | *Pocket* |
| $Y_1$ | Cβ | 4.0 | W167 | Cβ | A |
| | Cβ | 3.4 | W167 | Cγ | |
| | Cβ | 3.5 | W167 | Cδ1 | |
| | Cβ | 3.7 | W167 | Nε1 | |
| | Cβ | 3.8 | W167 | Cε2 | |
| | Cβ | 3.6 | W167 | Cδ2 | |
| | Cγ | 4.0 | W167 | Cδ1 | |
| | Cγ | 3.7 | W167 | Nε1 | |
| | Cγ | 3.6 | W167 | Cε2 | |
| | Cγ | 3.9 | W167 | Cδ2 | |
| | Cγ | 4.0 | W167 | Cζ2 | |
| | Cδ1 | 3.9 | Y60 | Cδ1 | |
| | Cδ1 | 3.6 | Y60 | Cε1 | |
| | Cδ1 | 3.7 | Q64 | Cγ | |
| | Cδ1 | 3.7 | Q64 | Cδ | |
| | Cε1 | 4.0 | Y60 | Cδ1 | |
| | Cε1 | 4.0 | Y60 | Cδ1 | |
| | Cε1 | 3.7 | Q64 | Cγ | |
| | Cε1 | 4.0 | Q64 | Cδ | |
| | Cε2 | 3.8 | W167 | Nε1 | |
| | Cε2 | 4.0 | W167 | Cε2 | |
| | Cε2 | 3.8 | W167 | Cζ2 | |
| | Cδ2 | 3.7 | W167 | Cδ1 | |
| | Cδ2 | 3.1 | W167 | Nε1 | |
| | Cδ2 | 3.4 | W167 | Cε2 | |
| | Cδ2 | 3.7 | W167 | Cζ2 | |
| | C | 3.9 | Y9 | Cζ | |
| | C | 4.0 | Y9 | Cε2 | |
| $M_2$ | Cα | 3.8 | Y9 | Cζ | B |
| | Cβ | 3.7 | Y9 | Cε1 | |
| | Cβ | 3.8 | Y9 | Cζ | |
| | Cγ | 4.0 | Y9 | Cε1 | |
| | Cγ | 4.0 | Y9 | Cδ1 | |
| | Sδ | 3.6 | A26 | Cβ | |
| | Sδ | 4.0 | M45 | Sδ | |
| | Sδ | 3.7 | M45 | Cε | |
| | Cε | 3.9 | M45 | Sδ | |
| | Cε | 3.5 | G68 | Cα | |
| $T_3$ | Cα | 3.9 | I67 | Cγ1 | - |
| | Cα | 4.0 | I67 | Cδ1 | |
| $L_4$ | Cβ | 3.9 | Y99 | Cε2 | D |

| | | | | | |
|---|---|---|---|---|---|
| | Cβ | 3.8 | Y159 | Cγ | |
| | Cβ | 4.0 | Y159 | Cδ1 | |
| | Cβ | 4.0 | Y159 | Cδ2 | |
| | Cδ1 | 3.7 | Y159 | Cβ | |
| | Cδ1 | 3.9 | Y159 | Cγ | |
| | Cδ1 | 3.8 | R155 | Cζ, A | |
| | Cδ1 | 3.8 | R155 | Cγ, A | |
| | Cδ2 | 3.8 | Y159 | Cδ2 | |
| | Cδ2 | 3.9 | Y99 | Cδ2 | |
| $Q_5$ | Cα | 3.6 | I67 | Cγ2 | - |
| | Cβ | 3.8 | I67 | Cγ2 | |
| | Cγ | 4.0 | G70 | C | |
| | Cδ | 3.8 | G70 | Cα | |
| | Cδ | 3.9 | G70 | C | |
| $A_6$ | Cα | 3.9 | R97 | Cζ | C |
| | Cβ | 3.9 | Y75 | Cε1 | |
| | Cβ | 3.5 | Y75 | Cζ | |
| | Cβ | 3.9 | Y75 | Cε2 | |
| | Cβ | 3.8 | V74 | Cγ1 | |
| | C | 3.7 | V74 | Cγ1 | |
| $V_7$ | Cβ | 3.9 | W146 | Cζ2 | E |
| | Cγ1 | 3.7 | W146 | Cε2 | |
| | Cγ1 | 3.9 | W146 | Cζ2 | |
| | C | 3.6 | V74 | Cγ1 | |
| | C | 4.0 | W146 | Cζ2 | |
| $T_8$ | Cα | 3.6 | V74 | Cγ1 | - |
| | Cβ | 3.9 | V74 | Cγ1 | |
| $F_9$ | Cγ | 3.8 | F122 | Cε2 | F |
| | Cδ1 | 3.8 | W146 | Cζ2 | |
| | Cδ1 | 3.7 | F122 | Cε2 | |
| | Cε1 | 3.9 | F115 | Cγ | |
| | Cε1 | 3.6 | F115 | Cδ2 | |
| | Cε1 | 3.8 | F115 | Cε2 | |
| | Cε1 | 4.0 | F122 | Cδ2 | |
| | Cε1 | 3.9 | F122 | Cε2 | |
| | Cζ | 3.8 | V95 | Cγ1 | |
| | Cζ | 3.9 | F115 | Cγ | |
| | Cζ | 3.4 | F115 | Cδ2 | |
| | Cζ | 3.9 | F115 | Cε2 | |
| | Cζ | 3.9 | F122 | Cδ2 | |
| | Cδ2 | 3.7 | S78 | Cβ | |
| | Cδ2 | 3.8 | L82 | Cγ | |
| | Cδ2 | 3.6 | L82 | Cδ1 | |
| | Cδ2 | 3.9 | F122 | Cε2 | |
| | Cε2 | 3.8 | V95 | Cγ1 | |
| | Cε2 | 3.8 | L82 | Cδ1 | |
| | Cε2 | 4.0 | F122 | Cε2 | |

**Table S3: Interactions of P3 within the Acar3 PBG**

| P3 | | | Acar3 | | |
|---|---|---|---|---|---|
| ***Hydrogen bonds (≤ 3.4 Å)*** | | | | | |
| *Residue* | *Atom* | *Distance (Å)* | *Residue* | *Atom* | *Pocket* |
| $M_1$ | O | 3.2 | Q64 | Nε2 | - |
| $T_2$ | Oγ1 | 2.7 | Y9 | Oη | A |
| | Oγ1 | 2.5 | Y171 | Oη | |
| | O | 2.5 | Y159 | Oη | |
| $M_3$ | - | - | - | - | B |
| $I_4$ | N | 3.1 | Y99 | Oη | D |
| | O | 3.0 | R97 | Nη1 | |
| | O | 3.2 | R97 | Nη2 | |
| $T_5$ | Oγ | 2.4 | R155 | Nη2 | - |
| | O | 2.8 | R155 | Nη1 | |
| | O | 3.2 | R155 | Nη2 | |
| $P_6$ | - | - | - | - | C |
| $P_7$ | - | - | - | - | E |
| $T_8$ | O | 3.0 | R145 | Nη2 | - |
| | O | 3.1 | W146 | Nε1 | |
| $F_9$ | N | 3.3 | S78 | Oγ | F |
| | O | 3.3 | R85 | Nη2 | |
| | O | 3.0 | T142 | Oγ1 | |
| | O | 2.6 | R145 | Nη1 | |
| ***Water-bridged hydrogen bonds (≤ 3.4 Å)*** | | | | | |
| *Residue (P3)* | *Atom* | *Distance (Å) P2-residue – $H_2O$* | *Distance (Å) $H_2O$ – Acar3-residue* | *Residue (Acar3)* | *Atom* |
| $P_7$ | O | 2.8 | 2.6 | S78 | Oγ |
| ***Salt bridges (≤ 4.0 Å)*** | | | | | |
| *Residue* | *Atom* | *Distance (Å)* | *Residue* | *Atom* | *Pocket* |
| $F_9$ | O | 3.3 | R85 | Nη2 | F |
| | | 2.6 | R145 | Nη1 | |
| | | 3.6 | | Nη2 | |
| ***Hydrophobic contacts/π-stacking (≤ 4.0 Å)*** | | | | | |
| *Residue* | *Atom* | *Distance (Å)* | *Residue* | *Atom* | *Pocket* |
| $M_1$ | Cβ | 4.0 | I67 | Cδ1 | - |

| | | | | | |
|---|---|---|---|---|---|
| | Cγ | 4.0 | I67 | Cδ1 | |
| $T_2$ | Cα | 3.8 | W167 | Cδ1 | A |
| | Cα | 4.0 | W167 | Cγ | |
| | Cβ | 3.7 | W167 | Cγ | |
| | Cβ | 4.0 | W167 | Cδ1 | |
| | Cβ | 3.9 | W167 | Cδ2 | |
| | Cγ2 | 3.2 | Q64 | Cδ | |
| | Cγ2 | 3.9 | Y60 | Cε1 | |
| | C | 3.9 | Y9 | Cε2 | |
| | C | 3.8 | Y159 | Cε1 | |
| $M_3$ | Cβ | 3.8 | I67 | Cγ2 | B |
| | Cγ | 3.9 | Y9 | Cγ | |
| | Cγ | 3.5 | Y9 | Cδ1 | |
| | Cγ | 3.5 | Y9 | Cε1 | |
| | Cγ | 3.9 | Y9 | Cζ | |
| | Sδ | 4.0 | Y9 | Cδ1 | |
| | Sδ | 4.0 | Y9 | Cε1 | |
| | Sδ | 3.7 | M45 | Cε | |
| | Sδ | 3.9 | A26 | Cβ | |
| | Cε | 3.5 | I67 | Cγ2 | |
| $I_4$ | Cγ1 | 3.7 | Y99 | Cε2 | D |
| | Cγ1 | 4.0 | Y159 | Cε2 | |
| | Cγ1 | 3.8 | Y159 | Cγ | |
| | Cγ1 | 3.9 | Y159 | Cδ2 | |
| | Cγ1 | 4.0 | Y159 | Cε1 | |
| | Cγ1 | 4.0 | Y159 | Cζ | |
| | Cγ1 | 3.9 | Y159 | Cδ1 | |
| | Cγ2 | 3.9 | Y159 | Cγ | |
| | Cγ2 | 3.5 | Y159 | Cδ1 | |
| | Cδ1 | 4.0 | Y159 | Cε2 | |
| | Cδ1 | 3.8 | Y159 | Cβ | |
| | Cδ1 | 3.5 | Y159 | Cγ | |
| | Cδ1 | 4.0 | Y159 | Cδ1 | |
| | Cδ1 | 3.4 | Y159 | Cδ2 | |
| | C | 4.0 | I67 | Cγ2 | |
| $T_5$ | - | - | - | - | - |
| $P_6$ | Cα | 3.6 | R97 | Cζ | C |
| | Cβ | 4.0 | F115 | Cε2 | |
| | Cβ | 3.9 | Y75 | Cε1 | |
| | Cβ | 3.8 | Y75 | Cζ | |
| | Cβ | 3.8 | R97 | Cζ | |
| | Cγ | 4.0 | V74 | Cβ | |
| | Cγ | 3.8 | V74 | Cγ1 | |
| | Cγ | 3.8 | Y75 | Cε1 | |
| | Cγ | 3.6 | Y75 | Cζ | |
| | Cγ | 3.8 | Y75 | Cε2 | |
| | Cδ | 4.0 | S71 | Cα | |

| | | | | | |
|---|---|---|---|---|---|
| | Cδ | 3.7 | S71 | Cβ | |
| $P_7$ | Cβ | 3.9 | W146 | Cε2 | E |
| | Cβ | 3.6 | V152 | Cγ1 | |
| | Cβ | 3.6 | W146 | Cζ2 | |
| | Cγ | 4.0 | W146 | Cζ2 | |
| | Cγ | 3.5 | V152 | Cγ1 | |
| | Cγ | 4.0 | Q156 | Cδ | |
| | Cδ | 3.6 | F115 | Cζ | |
| $T_8$ | Cα | 3.8 | V74 | Cγ1 | - |
| | Cβ | 4.0 | V74 | Cγ1 | |
| $F_9$ | Cγ | 3.9 | F122 | Cε2 | F |
| | Cδ1 | 3.9 | F112 | Cε2 | |
| | Cδ1 | 4.0 | T142 | Cγ2 | |
| | Cδ1 | 3.9 | W146 | Cζ2 | |
| | Cε1 | 4.0 | F122 | Cδ2 | |
| | Cε1 | 3.9 | F122 | Cε2 | |
| | Cε1 | 4.0 | W146 | Cη2 | |
| | Cε1 | 3.9 | F115 | Cδ2 | |
| | Cε1 | 4.0 | F115 | Cε2 | |
| | Cζ | 3.7 | F122 | Cδ2 | |
| | Cζ | 3.9 | F122 | Cε2 | |
| | Cζ | 4.0 | F115 | Cβ | |
| | Cζ | 3.7 | F115 | Cγ | |
| | Cζ | 3.4 | F115 | Cδ2 | |
| | Cζ | 3.9 | F115 | Cε2 | |
| | Cδ2 | 3.8 | L82 | Cγ | |
| | Cδ2 | 3.7 | L82 | Cδ1 | |
| | Cδ2 | 3.8 | L82 | Cδ2 | |
| | Cδ2 | 3.8 | F122 | Cε2 | |
| | Cδ2 | 3.6 | S78 | Cβ | |
| | Cε2 | 3.8 | L82 | Cδ1 | |
| | Cε2 | 3.8 | F122 | Cδ2 | |
| | Cε2 | 3.8 | F122 | Cε2 | |
| | C | 4.0 | R145 | Cζ | |

**Table S4: Comparison of r.m.s.d. values of anchor residues and terminal and central segments of peptides bound to Acar3, HLA-A*24:02 and HLA-B*15:01.**

| PDB code | Anchor residues/ positions | Peptide length (# of aa) | r.m.s.d. range/average (Å)* of: N-terminal anchor | aa 1-3** | aa 4-6** | aa 7-9** | C-terminal anchor |
|---|---|---|---|---|---|---|---|
| | | | ***Acar3*** | | | | |
| 7ZQI (P2) | M/2, F/9 | 9 | - /**0.87** | 0.87-1.31 /**1.09** | 0.64-1.60 /**1.01** | 0.96-1.22 /**1.07** | - /**1.03** |
| 7ZQJ (P3) | M/3, F/9 | 9 | | | | | |
| | | | ***HLA-A*24:02*** | | | | |
| 2BCK (A, B, C) | Y/2, L/9 | 9 | 0.15-0.58 /**0.35** | 0.48-0.54 /**0.52** | 1.29-3.16 /**2.27** | 0.87-1.32 /**1.10** | 0.11-0.95 /**0.66** |
| 3I6L | F/2, L/9 | 9 | | | | | |
| 4F7M (A, B, C) | Y/2, F/10 | 10 | | | | | |
| 4F7T (A, B, C) | Y/2, F/8 | 8 | | | | | |
| 5WWU | Y/2, F/11 | 11 | | | | | |
| 5WXD | Y/2, F/11 | 11 | | | | | |
| | | | ***HLA-B*15:01*** | | | | |
| 1XR8 | E/2, Y/9 | 9 | 0.02-0.72 /**0.48** | 0.02-1.48 /**0.50** | 0.45-3.57 /**1.75** | 0.12-1.29 /**0.52** | 0.12-1.16 /**0.59** |
| 1XR9 | L/2, Y/9 | 9 | | | | | |
| 3C9N | Q/2, M/9 | 9 | | | | | |
| 5TXS | Q/2, Y/9 | 9 | | | | | |
| 5VZ5 | Q/2, Y/10 | 10 | | | | | |

* The values were obtained using *Superpose* (Krissinel & Henrick, 2004) (*gesamt* method) from the *CCP4i program suite* (version 7.0.078) (Winn *et al.*, 2011) calculating r.m.s.d. values for all possible combinations of structures from the same protein. In HLA-A*24:02, conformational changes of F99 and Y159 may induce an expansion of pocket B. However, this mechanism rather enables the accommodation of a secondary anchor located at peptide position 3 [11] than providing additional flexibility to the primary anchor only. Consequently, those structures have been excluded from the r.m.s.d. calculations.

** The values each comprise the r.m.s.d. range and average of the three N-terminal, central and C-terminal residues combined from all structures containing nonameric peptides in all possible combinations. Due to the N-terminal variations of the peptides in the Acar3 structures, the superposition of P2 *vs.* P3 only contains two overlaps, *i.e.* residue 1 (P2) with residue 2 (P3) and residue 2 (P2) with residue 3 (P3).

**Table S5:** Half-life for peptide-MHC-I molecules for a series of peptides Acar3 and Acar19 combinations estimated using the dissociation using a scintillation proximity assay.

| Sequence | Stability T½ (h) | Acar |
|---|---|---|
| AANMYIYPL | 0,1 | Acar3 |
| FQFTLHWEL | 0,0 | Acar3 |
| KEISNQEPL | 0,0 | Acar3 |
| QQYAGWSAL | 0,0 | Acar3 |
| FLQGAKWYL | 0,1 | Acar3 |
| IMDASSFTL | 1,4 | Acar3 |
| KLMDVVYSI | 0,4 | Acar3 |
| SLAIDAYPL | 2,1 | Acar3 |
| AQRELFFTL | 0,0 | Acar3 |
| YLLADTFTV | 0,0 | Acar3 |
| VQYRGLYQI | 0,0 | Acar3 |
| RLHDAWWTL | 0,0 | Acar3 |
| SLAALFYSL | 0,0 | Acar3 |
| AMMWRIAQL | 0,3 | Acar3 |
| AELGAFFSI | 0,0 | Acar3 |
| MMHASTSPF | 4,2 | Acar3 |
| YLAENTFVV | 0,0 | Acar3 |
| KQLEWKWGI | 0,0 | Acar3 |
| WLAKSFFEL | 0,2 | Acar3 |
| SQEDNHFSL | 0,0 | Acar3 |
| AQIGIFAPV | 0,0 | Acar3 |
| KEMGFSPRL | 0,0 | Acar3 |
| SESTHYFTV | 0,0 | Acar3 |
| GLAATSFPL | 1,5 | Acar3 |
| GQWGGDWAV | 0,0 | Acar3 |
| SQDDNYFTL | 0,0 | Acar3 |
| LEMQHLISL | 0,0 | Acar3 |
| FQMGGIGPM | 0,7 | Acar3 |
| AQMVWIHGV | 0,0 | Acar3 |
| YMHGSIHEV | 0,9 | Acar3 |
| KQMEDGHTL | 0,0 | Acar3 |
| AEMRETHWL | 5,0 | Acar3 |
| SQAAFGLPI | 0,0 | Acar3 |
| KQLELFWVI | 0,2 | Acar3 |
| WQTDTTIPL | 0,0 | Acar3 |
| FLLENAAYL | 1,2 | Acar3 |
| WQLTSIWPI | 0,7 | Acar3 |
| NQFGSVPAL | 0,0 | Acar3 |
| AEAASATPL | 0,0 | Acar3 |
| FMIVNNFPV | 0,3 | Acar3 |
| RQWGMGFLL | 0,0 | Acar3 |
| LQAMHGFPL | 0,0 | Acar3 |
| MMQVWIQPL | 0,9 | Acar3 |
| TLKRRSWPL | 0,0 | Acar3 |

| | | |
|---|---|---|
| WMACNSAAF | 0,9 | Acar3 |
| KQSSFLSSL | 0,0 | Acar3 |
| YEFLGELAL | 0,0 | Acar3 |
| YLQQNTHTL | 0,5 | Acar3 |
| RQNAAIEAL | 0,6 | Acar3 |
| SQAFNTPAL | 0,0 | Acar3 |
| AELRHLNPL | 0,0 | Acar3 |
| YLACKQHAL | 0,0 | Acar3 |
| YQAGISAAL | 0,0 | Acar3 |
| SMFERDFHF | 0,2 | Acar3 |
| RLMKQDFSV | 0,0 | Acar3 |
| YLYLRPYAL | 2,4 | Acar3 |
| FQANMFLTL | 1,2 | Acar3 |
| REWSGLNYL | 1,9 | Acar3 |
| YQVKYVSPV | 3,6 | Acar3 |
| FQWWRSHPL | 0,0 | Acar3 |
| LEMCHSTQI | 0,0 | Acar3 |
| FLDKGTYTL | 0,3 | Acar3 |
| REAGMAATL | 0,0 | Acar3 |
| FMYSDFHFI | 0,1 | Acar3 |
| YLMPYSVYI | 0,1 | Acar3 |
| KQLESVMYL | 1,3 | Acar3 |
| ILMEHIHKL | 0,2 | Acar3 |
| KQWSWFSLL | 0,0 | Acar3 |
| LMDENTYAM | 0,4 | Acar3 |
| RQLLWRYQI | 0,0 | Acar3 |
| HELSLFWPL | 1,1 | Acar3 |
| MQQGRFPPL | 0,0 | Acar3 |
| RQQAIVDLL | 0,0 | Acar3 |
| RENGGYWLL | 0,0 | Acar3 |
| FMFDYIPPV | 0,2 | Acar3 |
| YQGSYGFRL | 0,0 | Acar3 |
| KQLEYSWVL | 0,1 | Acar3 |
| SQGRGWFLL | 0,0 | Acar3 |
| SMHFYGWSL | 0,2 | Acar3 |
| FQAQNIAGL | 0,0 | Acar3 |
| RMFKRVFNM | 0,0 | Acar3 |
| RQIRMTSTI | 0,1 | Acar3 |
| KAYANMWSL | 0,0 | Acar3 |
| RQYERYTAL | 0,0 | Acar3 |
| MQLPGGWLL | 0,0 | Acar3 |
| YQLGDYFFV | 0,0 | Acar3 |
| WEFVNRPPL | 0,0 | Acar3 |
| SQIETGTPF | 0,0 | Acar3 |
| AMPKTIYEL | 3,8 | Acar3 |

| | | |
|---|---|---|
| YLLEMLWRL | 0,4 | Acar3 |
| SQMGLSCAL | 0,3 | Acar3 |
| FQNWGIESI | 0,0 | Acar3 |
| REMINHYQV | 0,7 | Acar3 |
| AMAAGLSSL | 0,3 | Acar3 |
| DMSTNAEYF | 0,0 | Acar3 |
| GRWMLPQGM | 0,0 | Acar3 |
| KRKKAYADF | 0,0 | Acar3 |
| QAYAAPQLF | 1,5 | Acar3 |
| VQTAAAVVF | 0,2 | Acar3 |
| VMMAKRPLI | 0,5 | Acar3 |
| NTDEIPELI | 0,0 | Acar3 |
| IYLPIVHPF | 0,7 | Acar3 |
| GAMYALANF | 0,6 | Acar3 |
| YMKKRYEEF | 0,2 | Acar3 |
| AKIEAPLLL | 2,5 | Acar3 |
| WMMAMRYPI | 3,1 | Acar3 |
| KMNAKAATL | 2,1 | Acar3 |
| WRTIMAVLF | 0,0 | Acar3 |
| ERWFVRNPF | 0,0 | Acar3 |
| MTMITPPTF | 10,3 | Acar3 |
| QQMFPGAPF | 0,4 | Acar3 |
| AVMFFPFWF | 0,6 | Acar3 |
| TRAPAPFPL | 0,1 | Acar3 |
| GFMNEDHWF | 0,0 | Acar3 |
| DYQGLPLEF | 0,2 | Acar3 |
| EEMPLVWDL | 0,0 | Acar3 |
| KTAAAAAAF | 0,4 | Acar3 |
| YMWLGARYL | 0,4 | Acar3 |
| GMNVTAPAL | 0,3 | Acar3 |
| GRLVTVNPF | 0,5 | Acar3 |
| LDMNVVKEL | 0,6 | Acar3 |
| VTTPARTAF | 0,0 | Acar3 |
| MMMPMFNAF | 7,2 | Acar3 |
| YMTLQAVTF | 11,2 | Acar3 |
| VIMWYNYLF | 0,3 | Acar3 |
| SVYFAAFAF | 1,6 | Acar3 |
| ETTQALQLF | 0,0 | Acar3 |
| MPMSMPIPM | 3,2 | Acar3 |
| FRMLAWHVL | 0,3 | Acar3 |
| RRMATTFTF | 6,4 | Acar3 |
| QMSWLPLCV | 0,6 | Acar3 |
| LRQWAPATM | 0,0 | Acar3 |
| KAMRPWQSF | 0,4 | Acar3 |
| GTVPTDNPF | 0,0 | Acar3 |

| | | |
|---|---|---|
| LMMNGTSAM | 0,8 | Acar3 |
| QQLEADYTF | 0,4 | Acar3 |
| MTYKAAFDL | 1,8 | Acar3 |
| YYVWMVQFF | 0,0 | Acar3 |
| DIIRAHPWF | 0,0 | Acar3 |
| LPRWPPPQL | 0,3 | Acar3 |
| RMMETWHPL | 0,9 | Acar3 |
| FHMSKPSDL | 0,8 | Acar3 |
| LRWLAKSFF | 0,5 | Acar3 |
| KAMSTPFSL | 5,3 | Acar3 |
| LTMNLVSDI | 4,5 | Acar3 |
| KIMQVDRPM | 2,6 | Acar3 |
| SYPPPPASF | 0,2 | Acar3 |
| DTDPLPVVF | 0,0 | Acar3 |
| VTDTALAYF | 0,0 | Acar3 |
| QPWTPVSSF | 0,0 | Acar3 |
| VPMEKLKAL | 1,4 | Acar3 |
| LQMLGENVL | 2,6 | Acar3 |
| GMVGAEFSL | 7,6 | Acar3 |
| IRMEFQSVF | 0,4 | Acar3 |
| RMAMIPRTL | 0,5 | Acar3 |
| RMMATKDSF | 0,4 | Acar3 |
| RMLPKLAEF | 1,6 | Acar3 |
| VVKQLPASF | 0,1 | Acar3 |
| RMMGKTNPL | 0,2 | Acar3 |
| STEFIPNLF | 0,3 | Acar3 |
| VMPLLVWLF | 1,9 | Acar3 |
| SPMETTAEF | 3,0 | Acar3 |
| NMKWKFNAL | 0,0 | Acar3 |
| YMWECPDFF | 2,6 | Acar3 |
| LPLPAQQPF | 0,3 | Acar3 |
| KRLLLKLDF | 0,0 | Acar3 |
| WMYNIQPYL | 0,0 | Acar3 |
| RATQPAAEF | 1,6 | Acar3 |
| NMAPEKVDF | 0,0 | Acar3 |
| KMAAFPETL | 0,4 | Acar3 |
| ETMKPAAMV | 2,7 | Acar3 |
| YWMGGTTYF | 0,8 | Acar3 |
| RYMSKTYNF | 7,5 | Acar3 |
| GPMKLVMAF | 0,7 | Acar3 |
| EFKQILTDF | 0,0 | Acar3 |
| RRWIAPHPL | 1,7 | Acar3 |
| MMMSTAVAF | 7,3 | Acar3 |
| KRMMVRHCL | 0,0 | Acar3 |
| MMWATAQAL | 0,8 | Acar3 |

| | | |
|---|---|---|
| KTMMQAHDL | 9,9 | Acar3 |
| AMSAQAAAF | 11,3 | Acar3 |
| SMMPEAMTI | 0,9 | Acar3 |
| KQMSQPYAV | 4,0 | Acar3 |
| IFLKPEETF | 6,1 | Acar3 |
| VWKQLFPEL | 0,0 | Acar3 |
| YPMSIPATL | 0,7 | Acar3 |
| QWALLPYLF | 9,9 | Acar3 |
| DMWEHAFYL | 0,0 | Acar3 |
| DAAKNQVAM | 0,2 | Acar19 |
| TFMYVFSTF | 0,2 | Acar19 |
| NTDEIPELI | 0,2 | Acar19 |
| NTFAAWLPM | 0,6 | Acar19 |
| DPNPQEVVL | 0,4 | Acar19 |
| QYQYLSILF | 0 | Acar19 |
| LYGLITEQF | 0 | Acar19 |
| TYSPALNKM | 0,1 | Acar19 |
| MYLRTIYDL | 4 | Acar19 |
| QYPAFVLFI | 0,2 | Acar19 |
| ESFDLAGLF | 0 | Acar19 |
| VMKRNFIDF | 0,1 | Acar19 |
| IYASRPLDF | 0 | Acar19 |
| EYARQVQMI | 0,4 | Acar19 |
| MSADNAGAL | 0 | Acar19 |
| VSFSMVGLF | 0,1 | Acar19 |
| NPSVLKILL | 0,2 | Acar19 |
| TRAPAPFPL | 0,1 | Acar19 |
| DFIGKTIGF | 0 | Acar19 |
| YQAGISAAL | 0,2 | Acar19 |
| TYMFTHIDL | 0,1 | Acar19 |
| GVPELGAFF | 0 | Acar19 |
| SFHIIEWLF | 0,2 | Acar19 |
| NAFHHPSAV | 0,1 | Acar19 |
| SYLGLTQPF | 0,2 | Acar19 |
| SQAPLPCVL | 0 | Acar19 |
| IYPGVNHGF | 0 | Acar19 |
| DISPTNIPL | 0,2 | Acar19 |
| EGAGIDDPV | 0,2 | Acar19 |
| YPMSIPATL | 0,1 | Acar19 |
| GAAGARGAL | 0 | Acar19 |
| IASPAWFLF | 0,3 | Acar19 |
| ITAALAWSL | 0,1 | Acar19 |
| IYGIFQSTF | 0,1 | Acar19 |
| LANYAFFAI | 0 | Acar19 |
| DPKKTGGPI | 0 | Acar19 |

| | | |
|---|---|---|
| AYARNLDTL | 0 | Acar19 |
| DFIANTETI | 4,7 | Acar19 |
| DPLVIPFSF | 0 | Acar19 |
| RYFTVAFLF | 0 | Acar19 |
| GHFPLQHAL | 0,2 | Acar19 |
| HTAEIQQFF | 0,3 | Acar19 |
| IAARAVELL | 0 | Acar19 |
| RFNNLTVYF | 0 | Acar19 |
| ISASLAALF | 0,1 | Acar19 |
| DVVPMVTQM | 0,6 | Acar19 |
| YWARATVEL | 0 | Acar19 |
| RYSGFVRTL | 0 | Acar19 |
| EFVSANLAM | 0 | Acar19 |
| EDFEIFYNL | 0,4 | Acar19 |
| YTYPIAHTA | 0,1 | Acar19 |
| RYAYTSVEF | 0 | Acar19 |
| WDAYIPHYV | 0 | Acar19 |
| ITFHNQRDF | 0,2 | Acar19 |
| KYNSHHFTF | 0 | Acar19 |
| YISPIFIPM | 0,1 | Acar19 |
| DSPATLSAY | 0 | Acar19 |
| RYPGVMYAF | 0 | Acar19 |
| ITLPISASL | 0 | Acar19 |
| DYLVSTQEF | 0 | Acar19 |
| AFHQLVQVI | 0,3 | Acar19 |
| ITDWLNFTL | 0,1 | Acar19 |
| MYLRFAERL | 0,2 | Acar19 |
| IAKGIDAEF | 0,1 | Acar19 |
| DHIPIINTL | 0,3 | Acar19 |
| DMNLEQWSV | 0,4 | Acar19 |
| QFAGGSFDF | 0 | Acar19 |
| HAATNFREI | 0,1 | Acar19 |
| DAAVVFPPV | 10 | Acar19 |
| ATPYNPEDI | 0 | Acar19 |
| EFKQILTDF | 3,6 | Acar19 |
| LYIEILRLL | 0,1 | Acar19 |
| VFSPFGYSF | 0 | Acar19 |
| IFPSMYFLM | 0,1 | Acar19 |
| IQNATMDEF | 0,1 | Acar19 |
| KYAKFFQNF | 0 | Acar19 |
| FSIMIYFTF | 0,2 | Acar19 |
| DTRAIDQFF | 2,3 | Acar19 |
| IALPVAWLF | 0,1 | Acar19 |
| TTLSIYFLL | 0,1 | Acar19 |
| LRHSTAHLL | 0,1 | Acar19 |

| | | |
|---|---|---|
| KASSAWHYF | 0,2 | Acar19 |
| NYNGLLSSI | 19,1 | Acar19 |
| WYAPVALLF | 0,2 | Acar19 |
| RYPAIAGGI | 0 | Acar19 |
| LFPELDCFF | 1,6 | Acar19 |
| DSIMLTATF | 1,5 | Acar19 |
| LMNELGVPF | 0,6 | Acar19 |
| DTFGVIDTM | 0,2 | Acar19 |
| WYIKIFIII | 0 | Acar19 |
| FFKCIYRLF | 0 | Acar19 |
| DVKFHTQAF | 0 | Acar19 |
| IQNALEKAL | 0,1 | Acar19 |
| YSRMLYIEF | 0 | Acar19 |
| YTIDLNDAF | 0 | Acar19 |
| NMAPEKVDF | 0,2 | Acar19 |
| RYSNFAWYF | 0 | Acar19 |
| ISPRNYFTF | 0 | Acar19 |
| YYNAFQWAI | 0,4 | Acar19 |
| YQAYAAPQL | 0 | Acar19 |
| NPLEIYQEI | 0,8 | Acar19 |
| DVSPILEAY | 0,1 | Acar19 |
| KSNGAQQWL | 0,1 | Acar19 |
| RYWYFAAEL | 0 | Acar19 |
| DGFGVHLAF | 0,4 | Acar19 |
| NVGRILGYV | 0,5 | Acar19 |
| DALLNEENL | 2 | Acar19 |
| HYNAFHWAI | 0,3 | Acar19 |
| DQIRNMSVI | 0,3 | Acar19 |
| KSAQVPLPL | 0 | Acar19 |
| KDPENFDHF | 0,2 | Acar19 |
| LSDAIFDDL | 0 | Acar19 |
| NTARLMAGA | 0,2 | Acar19 |
| IPKYLEIEI | 0,2 | Acar19 |
| IAASNLEQF | 0,1 | Acar19 |
| NFGRVKVHF | 0,6 | Acar19 |
| DYQGLPLEF | 3,5 | Acar19 |
| EAAKNVECI | 0,4 | Acar19 |
| IASPILFPF | 0,5 | Acar19 |
| IPAPGLGAL | 0,2 | Acar19 |
| YYSNKVFPI | 0,2 | Acar19 |
| YRSMIAFKL | 0,1 | Acar19 |
| YYNAFHWAI | 0,4 | Acar19 |
| ENALLVALF | 0,1 | Acar19 |
| QRASNVFDL | 0,2 | Acar19 |
| EYAPFARLL | 1,1 | Acar19 |

| | | |
|---|---|---|
| VYFVLTDRF | 0 | Acar19 |
| MAAAAFPAL | 0 | Acar19 |
| DWMERIEDF | 1,4 | Acar19 |
| FFSPVIASL | 0,9 | Acar19 |
| HYANFHNYF | 0 | Acar19 |
| VYDPLQPEL | 0 | Acar19 |
| ISDPLTSGL | 0,1 | Acar19 |
| MFMMIFHFV | 0,4 | Acar19 |
| ISARGQELF | 0,1 | Acar19 |
| DTETLTTMF | 0,1 | Acar19 |
| YYKKTFSAL | 0 | Acar19 |
| EYKKFIATF | 0 | Acar19 |
| SYLQAIGIL | 0,1 | Acar19 |
| IQNKLSSTF | 0,2 | Acar19 |
| MPARLWLCL | 0,9 | Acar19 |
| IAAPNMIAV | 0,1 | Acar19 |
| VSARIAALF | 0 | Acar19 |
| KFHQIEKEF | 0,3 | Acar19 |
| MYYPAQLYL | 0 | Acar19 |
| DYMPSMKRF | 0 | Acar19 |
| MFGYNPYSL | 0 | Acar19 |
| HYNAFQWAI | 0,3 | Acar19 |
| LAASVFFTF | 0,2 | Acar19 |
| FWPQHFGLI | 0,1 | Acar19 |
| ITNPFFYQM | 0 | Acar19 |
| EYLRIPLTI | 0 | Acar19 |
| MYPFIFFIV | 0 | Acar19 |
| DFISMYFPW | 0,5 | Acar19 |
| YFSGIMVRL | 0,4 | Acar19 |
| DMWEHAFYL | 0,3 | Acar19 |
| KYAEAFQMV | 0,3 | Acar19 |
| RMAMIPRTL | 0 | Acar19 |
| NYPASLHKF | 0,3 | Acar19 |
| DSFPNSVTI | 1,4 | Acar19 |
| DVAKIEAPL | 11,4 | Acar19 |
| YTMELCGAM | 0,1 | Acar19 |
| GFNKLRSTL | 0,4 | Acar19 |
| VTNRHEEKF | 0,1 | Acar19 |
| LFQPLHTVM | 0 | Acar19 |
| SHAAIGAYL | 0 | Acar19 |
| ISGPIKHPL | 0 | Acar19 |
| STAPTGSWF | 0,2 | Acar19 |
| EAPENAREL | 0 | Acar19 |
| DAMENPLSL | 0,2 | Acar19 |
| KSAQFPFHF | 0 | Acar19 |

| | | |
|---|---|---|
| GYAGTLQSL | 0 | Acar19 |
| IFDDLQGSL | 0,1 | Acar19 |
| RPAPATGAL | 0,3 | Acar19 |
| GFPSLESSF | 0,1 | Acar19 |
| QWALLPYLF | 0 | Acar19 |
| SYSGQRTEL | 0 | Acar19 |
| YFDPANGKF | 0,1 | Acar19 |
| IPNYLEIEI | 0,6 | Acar19 |
| TFMGMNVQF | 0,1 | Acar19 |
| ITPTIYLLL | 0 | Acar19 |
| RYMYIYLFI | 0 | Acar19 |
| AYAGLFTPL | 0,4 | Acar19 |
| DYKECEWPL | 17,5 | Acar19 |
| QYSGFVRTL | 0 | Acar19 |
| IYNTVVLTI | 0 | Acar19 |
| SFIPIFYQF | 1,2 | Acar19 |
| GYAWIDFDI | 0,2 | Acar19 |
| IYKNKISDF | 0 | Acar19 |
| IFFFLFNIL | 0 | Acar19 |
| DVSPLMHLF | 2,9 | Acar19 |
| YFNTHDVYF | 0 | Acar19 |
| DFGYATMAK | 0,3 | Acar19 |
| DAVRNAAAI | 0,3 | Acar19 |
| SYAAAQRKL | 0 | Acar19 |
| IYYWTAWLI | 0,2 | Acar19 |
| NNKSRLVAF | 0 | Acar19 |
| RMPPLGHEL | 0 | Acar19 |
| FSPRLCRPF | 0,4 | Acar19 |
| NHIRIEYII | 0,4 | Acar19 |
| AYPELACAV | 0 | Acar19 |
| IYLRRYFEI | 0 | Acar19 |
| RYVGLYLPF | 0 | Acar19 |
| IYWLIFWRF | 0 | Acar19 |
| SYQALAFDI | 0,5 | Acar19 |
| ETFRLLRSF | 0,1 | Acar19 |
| FYPPVLQPI | 0,3 | Acar19 |
| FFSPFFFSL | 0,2 | Acar19 |
| LANPTADDF | 0 | Acar19 |
| VYAPAGVEL | 0 | Acar19 |
| WFGHLASDW | 0,1 | Acar19 |
| IYLPIVHPF | 0 | Acar19 |
| RSFRIHILF | 0 | Acar19 |
| TTKPFWFQL | 0 | Acar19 |
| SSNPVMSRF | 0,1 | Acar19 |
| TYLALMATF | 0,1 | Acar19 |

| | | |
|---|---|---|
| NYFNRMFHF | 0,7 | Acar19 |
| YYYNFSEDL | 0 | Acar19 |
| QTSSIEGAW | 0,1 | Acar19 |
| MGAGLVFPI | 0 | Acar19 |
| DYAMHGTVF | 2,4 | Acar19 |
| LYPTFYCLF | 0,4 | Acar19 |
| STYGISEDL | 0,1 | Acar19 |
| DVSEQLELI | 0 | Acar19 |
| NRYGLPEKM | 0 | Acar19 |
| DIQKLVGKL | 0,4 | Acar19 |
| ENARLRALL | 0 | Acar19 |
| DASFNASAI | 1 | Acar19 |
| DMSTNAEYF | 0,2 | Acar19 |
| DYLTTYFTW | 0 | Acar19 |
| NFWLNTLLF | 3,4 | Acar19 |
| MYMALIAAF | 0,3 | Acar19 |
| DFNEAIQAY | 0,3 | Acar19 |
| MYQYIFLSF | 0 | Acar19 |
| MAMGILHTI | 0,1 | Acar19 |
| LSNFMLWQF | 0 | Acar19 |
| LYNTAIVGL | 0 | Acar19 |
| QFLSFASLF | 0 | Acar19 |
| IYAAISYMI | 0,3 | Acar19 |
| RFNAIWFNH | 0 | Acar19 |
| RAAPLMQSL | 0 | Acar19 |
| MSNPLTSPI | 6,3 | Acar19 |
| VQHNIKHSF | 0,1 | Acar19 |
| EWAENCYNL | 0 | Acar19 |
| ISKKAKGWF | 0 | Acar19 |
| TATPAWDAL | 0,1 | Acar19 |
| TVADIWHAM | 0,1 | Acar19 |
| NYMPYVFTL | 14,9 | Acar19 |
| AFSPNRFWM | 0 | Acar19 |
| DPSGAYFAW | 0,1 | Acar19 |
| DWSGYSGSF | 0,3 | Acar19 |
| STFAASGPF | 0 | Acar19 |
| YNAELLVAL | 0,2 | Acar19 |
| DYLQYVLQI | 0,5 | Acar19 |
| HYQTLCTNF | 0 | Acar19 |
| IYTVIYYIF | 0,2 | Acar19 |
| TTNGCSQAM | 0 | Acar19 |
| ITGQIIFGF | 0 | Acar19 |
| ISPGNKSDM | 0 | Acar19 |
| IANTTDHFF | 0,2 | Acar19 |
| TNTQNNDWF | 0,2 | Acar19 |

| | | |
|---|---|---|
| AFHGLDVKF | 0 | Acar19 |
| IQNEIMKPL | 0,2 | Acar19 |
| DTAKAFAYL | 0,5 | Acar19 |
| YVFAIPLPF | 0 | Acar19 |
| DYLQCVLQI | 6,6 | Acar19 |
| DIKLIDIAL | 3,7 | Acar19 |
| DPNPQEMVL | 0,1 | Acar19 |
| ISSGCTKTF | 0,1 | Acar19 |
| ESPINQIAF | 0 | Acar19 |
| DTDPLPVVF | 0,2 | Acar19 |
| DYDDVVHEV | 0,2 | Acar19 |
| ITPKTHFLF | 0 | Acar19 |
| PYPYLFYKF | 0 | Acar19 |
| EAAGIAACL | 0,6 | Acar19 |

Consortium, T. U. (2020). UniProt: the universal protein knowledgebase in 2021. *Nucleic acids research, 49*(D1), D480-D489. doi:10.1093/nar/gkaa1100

Krissinel, E., & Henrick, K. (2004). Secondary-structure matching (SSM), a new tool for fast protein structure alignment in three dimensions. *Acta Crystallogr D Biol Crystallogr, 60*(Pt 12 Pt 1), 2256-2268. doi:10.1107/s0907444904026460

Rasmussen, M., Harndahl, M., Stryhn, A., Boucherma, R., Nielsen, L. L., Lemonnier, F. A., . . . Buus, S. (2014). Uncovering the peptide-binding specificities of HLA-C: a general strategy to determine the specificity of any MHC class I molecule. *J Immunol, 193*(10), 4790-4802. doi:10.4049/jimmunol.1401689

Saper, M. A., Bjorkman, P. J., & Wiley, D. C. (1991). Refined structure of the human histocompatibility antigen HLA-A2 at 2.6 Å resolution. *Journal of Molecular Biology, 219*(2), 277-319. doi:https://doi.org/10.1016/0022-2836(91)90567-P

Thomsen, M. C. F., & Nielsen, M. (2012). Seq2Logo: a method for construction and visualization of amino acid binding motifs and sequence profiles including sequence weighting, pseudo counts and two-sided representation of amino acid enrichment and depletion. *Nucleic acids research, 40*(Web Server issue), W281-W287. doi:10.1093/nar/gks469

Winn, M. D., Ballard, C. C., Cowtan, K. D., Dodson, E. J., Emsley, P., Evans, P. R., . . . Wilson, K. S. (2011). Overview of the CCP4 suite and current developments. *Acta Crystallogr D Biol Crystallogr, 67*(Pt 4), 235-242. doi:10.1107/s0907444910045749
